# Stealth CAR-Tregs evade CD3 antibody therapy and skew immune responses toward regulation

**DOI:** 10.64898/2026.10.13.758862

**Authors:** Tifanie Blein, Nicolas Ayas, Soëli Charbonnier, Artur Gil, Aness Haddouche, Jehanne Hassan, Lucas Rabaux, Isabelle André, Hermann Waldmann, Baptiste Lamarthée, Makoto Miyara, Lucienne Chatenoud, Juliette Leon, Julien Zuber

**Affiliations:** Institut Necker Enfants Malades, CNRS UMR8253, INSERM UMR1151, Paris, France; Institut Imagine, INSERM UMR 1163, Paris Descartes Université-Sorbonne Paris Cité, Paris, France; Université Paris Cité, Paris, France; Centre d’Immunologie et des Maladies Infectieuses, INSERM U1135, Hôpital Pitié-Salpêtrière AP-HP, Sorbonne Université, Paris, France; Sir William Dunn School of Pathology, Oxford, United Kingdom; Université Marie et Louis Pasteur, EFS-BFC, INSERM UMR RIGHT, Besançon, France; Assistance Publique–Hôpitaux de Paris, Hôpital Necker, Service des Maladies du Rein et du Métabolisme, Transplantation et Immunologie Clinique, Paris, France

**Keywords:** tolerance, transplantation, CAR-Treg, CD3 antibody, depletion

## Abstract

Organ transplantation standard of care relies on lifelong immunosuppression, increasing infection and malignancy risks. An emerging approach using Chimeric Antigen Receptor Regulatory T cells (CAR-Tregs) targeting donor antigens, allows the suppression of alloimmune responses while preserving protective immunity. However, transplant recipients harbor a substantial pool of alloreactive memory T-cells, limiting CAR-Tregs tolerance-promoting effect. To address this challenge, our strategy combines selective depletion of activated T-cells, using non-mitogenic CD3 antibody therapy, with HLA-A2-targeted-CAR-Treg therapy. We established that T-cell receptor alpha chain (*TRAC*) deletion does not impair CAR-Treg antigen-specific activation, expansion, or their transcriptomic, phenotypic, epigenetic, and functional profiles. *TRAC*-/-CAR-Tregs exhibit robust protection from CD3 antibody depletion in circulation and peripheral tissues, whereas most *TRAC*+/+CAR-T cells are eliminated. Importantly, *TRAC*+/+CAR-T cells that escape CD3 antibody mediated depletion exhibit a transcriptomic signature consistent with T-cell exhaustion. Our findings demonstrate that CD3 antibody therapy provides an *in vivo* survival advantage to *TRAC*-/-CAR-Tregs, thereby shifting the immune balance toward regulation. This approach might enhance the translational potential of CAR-Treg therapy by reducing the required cell dose needed for clinical efficacy.

## INTRODUCTION

Standard immunosuppressive therapy remains the cornerstone of care in organ transplantation, effectively preventing rejection. However, it requires lifelong administration and is associated with an increased risk of infections and malignancies. In contrast, donor-specific (ds-) regulatory T cell (Treg) therapy has emerged as a promising alternative, offering greater antigen specificity than conventional approaches based on polyclonal Tregs^1^. This strategy aims to selectively suppress alloimmune responses while preserving protective immunity.

Despite its potential, clinical trials in kidney and liver transplantation have underscored the challenges of generating sufficient numbers of ds-Tregs by co-culturing recipient-derived Tregs with donor antigen-presenting cells^2,3^. To overcome these limitations, chimeric antigen receptor (CAR) technology has been employed to redirect Tregs toward donor-specific antigens^4,5^. Recent comparative studies demonstrate that CAR-Tregs exhibit superior donor-specific suppressive function compared to ds-Tregs^6^. Two ongoing clinical trials are currently evaluating HLA-A2-targeted CAR-Tregs in kidney (NCT04817774) and liver transplantation (NCT05234190)^7^. Preliminary data from the LIBERATE study suggest that CAR-Tregs may enable substantial reduction of immunosuppression, consistently achieving a calcineurin inhibitor-free monotherapy. However, rejection episodes were frequently observed when complete withdrawal of immunosuppressive drugs was attempted. These findings suggest that, while CAR-Tregs may support immunosuppression minimization, they are insufficient to induce full operational tolerance in solid organ transplantation. This partial efficacy underscores the persistent challenge of tipping the immune balance toward regulation in transplantation, particularly given the high precursor frequency of alloreactive T cells^8^. Moreover, transplant recipients often harbor a large pool of alloreactive memory T cells, shaped by prior heterologous immune responses ^9–13^, which are relatively resistant to Treg-mediated suppression. Collectively, these observations highlight the need for complementary strategies that also target effector T cells. Supporting this concept, the most favorable outcomes reported to date in liver transplantation were achieved using a combinatorial approach that included lymphodepletion with cyclophosphamide prior to ds-Treg infusion^7,14^.

Building on this rationale, we developed a strategy that combines depletion of alloreactive T cells with the administration of HLA-A2-targeted CAR-Treg therapy. We selected the CD3 monoclonal antibody otelixizumab (CD3 Ab) as a debulking agent due to three key advantages. First, it preferentially targets activated and proliferating T cells, thereby reducing preferentially alloreactive clones while sparing quiescent T cells^15,16^; second, it promotes a TGF-β-rich, anti-inflammatory environment that supports Treg induction and function^17,18^. Third, teplizumab, another second-generation non-mitogenic CD3 Ab has been developed clinically for type 1 diabetes. Its use was approved by the FDA in 2022 and by the EMA in 2025.

A potential limitation of this approach, however, is the susceptibility of CAR-Tregs to CD3 Ab-mediated depletion. To circumvent this, we hypothesized that CRISPR-Cas9-mediated disruption of the T-cell receptor alpha chain (*TRAC*) gene would destabilize the endogenous TCR/CD3 complex, rendering CAR-Tregs resistant to anti-CD3-induced cytotoxicity while preserving their antigen-specific suppressive function.

## RESULTS

### TRAC-deficient CAR-Tregs lack surface expression of the TCR/CD3 complex

We first sought to determine whether disruption of the *TRAC* gene would lead to the complete dismantling of the TCR/CD3 complex without substantially affecting Treg biology. To this end, naïve Tregs and conventional T cells (Tconvs) were FACS-sorted from HLA-A2/A28-negative donors, activated via CD3/CD28 stimulation, and transduced with a lentiviral vector (LV) encoding an HLA-A2-targeted CAR **(supplementary figure 1A)**. Two days post-transduction, the *TRAC* gene was disrupted using CRISPR-Cas9-mediated gene editing. Transduced T cells expressing EGFRt were sorted by FACS **(supplementary figure 1B)** and subsequently restimulated through CAR engagement on day 8, prior to further *in vitro* analysis on days 10 and 15 **(figure 1A)**.

**Figure 1:**
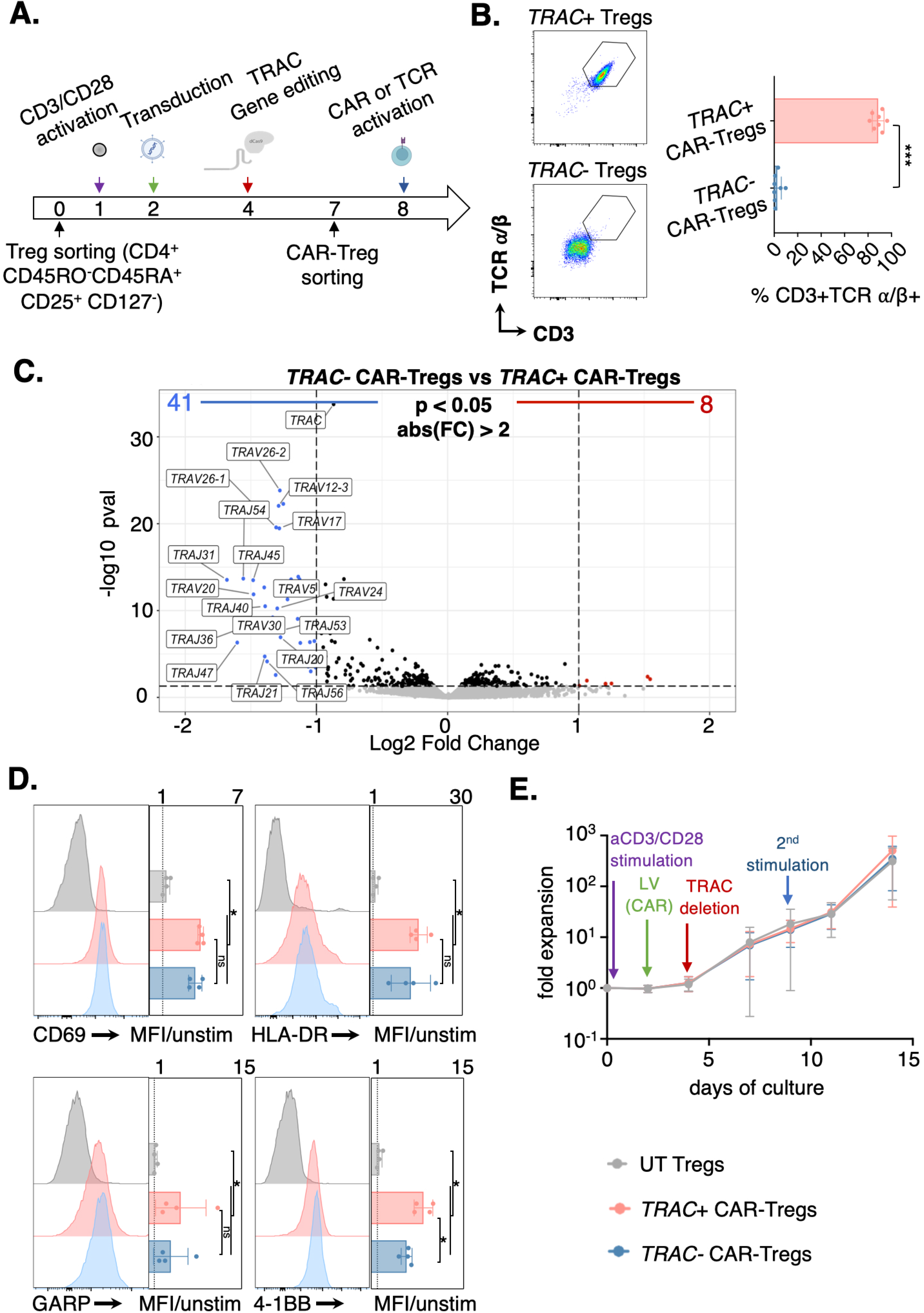
*TRAC*-deficient CAR-Tregs keep functional responses and expansion upon HLA-A2 specific CAR engagement, independently of endogenous TCR expression. **A.** Process timeline for *TRAC*-deficient CAR-Treg production. Naïve Tregs were isolated from healthy donor peripheral blood, polyclonally activated and subsequently transduced with the CAR and *TRAC* gene edited. CAR-Tregs were re-stimulated through the CAR before phenotypic, transcriptomic, epigenetic, and functional analyses. UT Tregs were restimulated polyclonally. **B.** CD3 and TCRα/β co-expression frequency, as assessed by flow cytometry at day 7, is compared across gene edited cells with either *TRAC*-targeted gRNA or scramble gRNA with a two-tailed Mann–Whitney test. *TRAC*-edited cells (*TRAC-/-* CAR-Tregs) lost both CD3 and TCRα/β surface expression: ***p<0.0002 (Mean ± SD). n= 8 donors over 8 independent experiments. **C.** Volcano plot showing differential gene expression between *TRAC-/-* CAR-Tregs vs. *TRAC*+/+ CAR-Tregs (log2 FC vs. − log10 p-value). Transcriptomic data were obtained from n=4 independent experiments, with live FACS-sorted cells on day 15 of culture. **D.** At day 10, cells were restimulated with HLA-A2-expressing B cells and activation marker expression was assessed by flow cytometry after 40 hours. Representative histograms (left) and MFI ratio (fold change compared to unstimulated controls) (right) are shown. Data from n=4 donors from 4 independent experiments. The mean ± SD is represented. Two-tailed Mann–Whitney tests were performed to compare CAR-Tregs (either *TRAC-/-* or *TRAC+/+*) with UT-Tregs *p < 0.05. Statistically significant induction of CD69, HLA-DR, GARP and 4-1BB markers was observed (p=0.0286). **E.** Fold expansion of UT Tregs and CAR-Tregs (either *TRAC-/-* or *TRAC+/+*) over a 14-day culture period. The mean ± SD is shown. A total of n=10 independent experiments are shown including the following cell populations: UT Tregs (grey full circle), *TRAC+/+* CAR-Tregs (red full circle) and *TRAC-/-* CAR-Tregs (blue full circle). Two-tailed Mann–Whitney indicated no significant differences in expansion between UT and CAR-Tregs at any timepoints.

Given that the TCR alpha chain, particularly its transmembrane domain, is essential for maintaining the structural integrity of the TCR/CD3 complex^19^, we anticipated complete destabilization of the complex following *TRAC* disruption. Consistent with this hypothesis, flow cytometry analysis revealed that surface expression of the TCRα/β and CD3 was nearly abrogated by day 3 post-editing in *TRAC*-targeted cells, with a mean knockout efficiency exceeding 97.4%±3.44, in stark contrast to unedited controls **(figure 1B)**.

To assess whether the absence of endogenous TCR tonic signaling leads to significant alterations in Treg biology, bulk RNA-sequencing was performed on *TRAC*-sufficient and - deficient CAR-Tregs on day 15 of culture (**figure 1C and supplementary table 1 and supplementary figure 1C**). Transcriptomic analysis indicated that gene editing primarily affected *TRAC*-related genes, with minimal impact on other transcripts. Overall, the two CAR-Treg populations displayed highly similar gene expression profiles, with only a small subset of differentially expressed genes. Notably, these included genes encoding for T cell receptor alpha variable (TRAV) and joining (TRAJ) segments involved in TCR rearrangement.

### CAR-Tregs exhibit functional responses upon HLA-A2 specific CAR engagement, independent of endogenous TCR expression

To evaluate whether CAR-mediated activation of *TRAC*-deficient Tregs is influenced by the absence of the endogenous TCR/CD3 complex, CAR-Tregs were co-cultured overnight with HLA-A2+ splenic B cells. Notably, CAR engagement induced the upregulation of CD69, HLA-DR, GARP and 4-1BB regardless of TCR expression, with a slightly higher 4-1BB levels in TCR-expressing cells. This indicates that CAR signaling alone is sufficient to trigger Treg activation. In contrast, UT Treg controls did not become activated upon exposure to HLA-A2-expressing stimulators (**figure 1D**).

Given the current challenge of generating sufficient numbers of ds-Tregs to achieve therapeutic efficacy, we next investigated whether deletion of the TCR complex affects the proliferative capacity of CAR-Tregs. No significant difference in fold expansion were observed over a 14-days culture period among UT Tregs, *TRAC*-sufficient CAR-Tregs, and *TRAC*-deficient CAR-Tregs following stimulation with anti-CD3/CD28 and HLA-A2, respectively (**figure 1E**). Optimization of the CAR-Treg manufacturing protocol consistently exceeded a 350-fold expansion in both CAR-Tregs populations over the 14-days culture period.

Importantly, compared with the original protocol described earlier^5^, fold expansion at day 7 after the second stimulation was markedly higher, reaching 153.3±104.7 for *TRAC*-sufficient CAR-Tregs and 104.9±60.3 for *TRAC*-deficient CAR-Tregs, versus 35.44±46.3 for *TRAC*-sufficient CAR-Tregs generated using the initial regimen (**supplementary figure 2A**).

### TRAC-deficient CAR-Tregs preserve core regulatory T cell transcriptomic identity

Forkhead box 3 (FOXP3) expression, a hallmark of Treg lineage commitment, is tightly controlled by transcription factors downstream of TCR signaling. These factors act via binding to the *FOXP3* promoter and conserved non-coding regulatory elements (CNS). Given FOXP3’s pivotal role in maintaining Treg stability and function, we investigated whether signaling through a CD28-CD3ζ CAR is sufficient to preserve Treg identity in the absence of endogenous TCR signaling. Notably, the frequency of FOXP3-expressing CAR-Treg remained high, 90.9%±4.2 and 90.7%±4.3 on day 11 and 83.0%±10.4 and 83.9%±9.4 on day 14 of culture, for *TRAC*-sufficient and -deficient CAR Tregs respectively. The majority of these cells retained HELIOS expression, regardless of *TRAC* expression status (**supplementary figure 2B**). In contrast, Tconvs exhibited negligible frequencies of FOXP3 and HELIOS co-expression (**figure 2A**). Moreover, the proportion of FOXP3-HELIOS-double-negative cells, indicative of Treg destabilization, remained low across all Treg populations (**supplementary figure 2B**) throughout the culture period.

**Figure 2:**
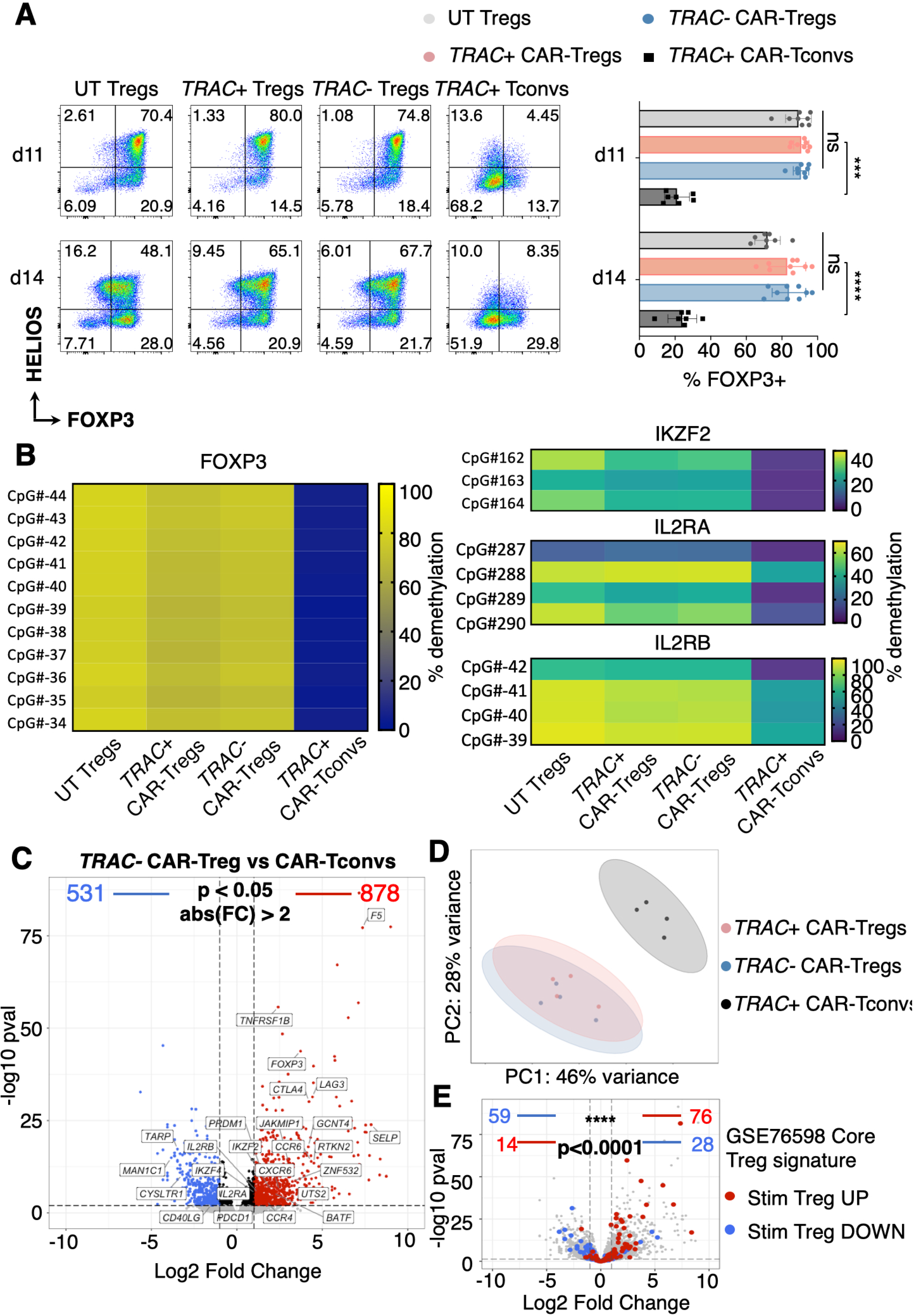
*TRAC*-deficient CAR-Tregs preserve core phenotypic, transcriptomic and epigenetic regulatory T cell identity. **A.** Flow-cytometry analysis of HELIOS and FOXP3 expression. (Left panel) Representative dot plots on day 11 and 14 of culture. (Right panel) Corresponding frequencies of FOXP3-positive cells from at least n=8 independent experiments. The mean ± SD is shown. All Tregs populations express significantly more FOXP3 on days 11 (p=0.0003) and 14 (p<0.0001). **B.** Bisulfite-conversion-based methylation PCR analysis on day 15 showing the percent demethylation of (left panel) the Treg-specific demethylated region (TSDR) and (right panel) IKZF2 (HELIOS), IL2RA (CD25) and IL2RB (CD122) of UT Tregs, *TRAC-/-* CAR-Tregs, *TRAC+/+* CAR-Tregs and CAR-Tconvs from n=4 independent experiments. **C-E.** Transcriptomic data from n=4 independent experiments, with live FACS-sorted cells (*TRAC-/-* CAR-Tregs, *TRAC+/+* CAR-Tregs and CAR-Tconvs) on day 15 of culture. **C.** Volcano plot showing differential gene expression between *TRAC-/- CAR*-Tregs vs. *TRAC+/+ CAR*-Tconvs (log2 FC vs. − log10 p-value). **D**. Principal component analysis (PCA) representing the *TRAC-/-* CAR-Tregs, *TRAC+/+* CAR-Tregs and CAR-Tconvs based on the top 2 most significantly contributing components of transcriptomic data. **E.** A Treg gene signature from^22^ is mapped on the *TRAC*-/- CAR-Tregs vs. *TRAC*+/+ CAR-Tconvs volcano plot (same as **C**). Values shown at the top represent the number of genes upregulated (red) or downregulated (blue) in both the Treg signature and our dataset. The second row indicates the number of genes upregulated (red) or downregulated (blue) in the Treg signature but regulated in the opposite direction in our dataset. Statistical significance of signature enrichment was assessed using a χ² test. ****p < 0.0001.

To further assess lineage stability in *TRAC*-deficient Tregs, we analyzed the DNA methylation profiles of key regulatory loci associated with Treg identity. The Treg-specific demethylated region (TSDR) within the CNS2 of the *FOXP3* gene serves as a robust epigenetic marker of stable Tregs^20,21^. All CpG sites within the *FOXP3* locus remained equivalently hypomethylated in both *TRAC*-sufficient (68.2%±13.6) and -deficient CAR-Tregs (71.8%±12.2). Additionally, key Treg-associated genes, including *IKZF2*, *IL2RA* and *IL2RB* maintained high levels of hypomethylation relative to Tconvs (**figure 2B**).

Crucially, the methylation status of core regulatory loci remained consistent with a stable Treg phenotype (**supplementary figure 2C**). Collectively, these findings demonstrate that CAR signaling via CD28-CD3ζ is sufficient to sustain FOXP3 expression and preserve the epigenetic integrity of Tregs, even in the absence of endogenous TCR expression.

To further assess the impact of TCR ablation on Treg identity, we compared CAR-restimulated *TRAC-*sufficient and -deficient CAR-Tregs global transcriptomic profiles with those of CAR-Tconvs. Consistent with the observation that TCR deficiency does not induce major transcriptomic changes (**figure 1C**), both *TRAC*-sufficient and -deficient CAR-Tregs maintained expression of Treg-lineage hallmark genes that distinguish them from CAR-Tconvs (**figures 2C and 2D**).

These findings indicate that *TRAC*-deficient CAR-Tregs retain a canonical Treg transcriptomic signature, including sustained expression of genes associated with lineage stability (*FOXP3, IKZF2*), suppressive function (*CTLA-4, LAG3, IL10, FCRL3* and *ARG1*), activation (*TNFRSF9* and *TNFRSF1B*) and migration (*CXCR6* and *CCR6*). Importantly, the *TRAC-*deficient CAR-Treg population conserved a well-defined Treg signature^22^, established through comparison of stimulated Tregs and stimulated Tconvs (**figures 2C and 2E**).

### CAR-Tregs exert a distinct subset with an effector Treg profile upon CAR stimulation

To capture potential post-translational changes induced by TCR deletion, we next assessed the expression of proteins associated with Treg identity, activation, functionality, trafficking and suppression (**supplementary table 2**) using high dimensional mass cytometry by time-of-flight (CyTOF). CAR-Treg and CAR-Tconv populations were analyzed 7 days following CAR stimulation. Similar to what was observed at the transcriptomic level, both CAR-Treg populations clustered together independently of TCR expression, and were clearly separated from CAR-Tconvs (**figure 3A and 3B, and supplementary figure 3)**. While maintaining high expression of key Treg lineage features (FOXP3, CD25, and HELIOS), both CAR-Treg populations contained a distinct cluster of activated effector Treg cells co-expressing HLA-DR, CD71, ICOS, TIGIT, OX40, TOX, PD-1, TIM-3, and CTLA-4 (**figure 3B**). Taken together, these results corroborate the transcriptomic data (**Supplementary figure 4**) and indicate the emergence of an effector Treg phenotype following CAR-specific stimulation, independent of endogenous TCR expression.

**Figure 3:**
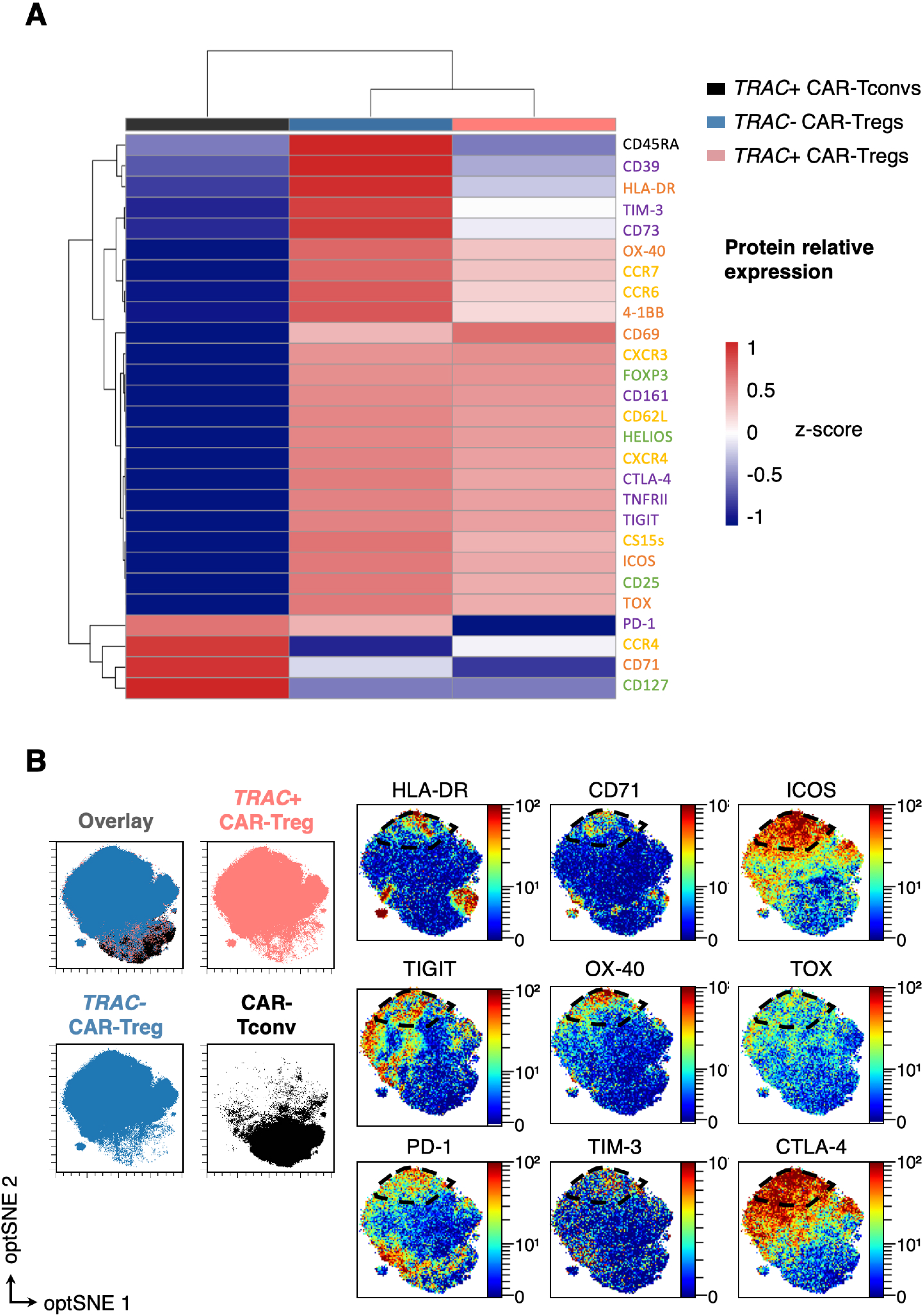
Both *TRAC*-/- and *TRAC*+/+ CAR-Tregs keep key Tregs protein expression and contain a distinct subset with effector Treg profile upon CAR stimulation. **A-B.** Protein expression of key Treg markers assessed by high-dimensional phenotypic analysis by Cytometry by Time-Of-Flight (CyTOF, mass cytometry) on day 15 of culture. Merged samples of n=3 different donors. **A.** Heatmap showing the median protein expression of markers related to Treg identity (green), trafficking (yellow), activation and functionality (orange), and suppression (purple) for each cell population, gated on total viable cells. Euclidean hierarchical clustering was applied, and protein expression values were normalized using z-scores. **B.** (Left panel) Spatial distribution of each cell population visualized by 2D opt-SNE: *TRAC*+/+ CAR-Tregs (red), *TRAC*-/- CAR-Tregs (blue), and CAR-Tconvs (black). (Right panel) Protein expression of key effector Treg markers projected onto the same 2D opt-SNE map.

### TRAC-deficient CAR-Tregs are as suppressive as TRAC-sufficient CAR-Tregs

We previously demonstrated that *TRAC*-deficient CAR-Tregs express CTLA-4 at levels comparable to *TRAC*-sufficient CAR-Treg (**figures 3 and 4A**). CTLA-4 is a critical mediator of Treg function, enabling the removal of costimulatory molecules CD80 and CD86 from the surface of antigen-presenting cells (APCs) via transendocytosis (**figure 4B**)^23^. This mechanism of APC suppression, which can be robustly quantified *in vitro*, has been shown to correlate with the *in vivo* suppressive efficacy of CAR-Tregs in a xenogeneic graft-vs-host disease model^4^.

**Figure 4:**
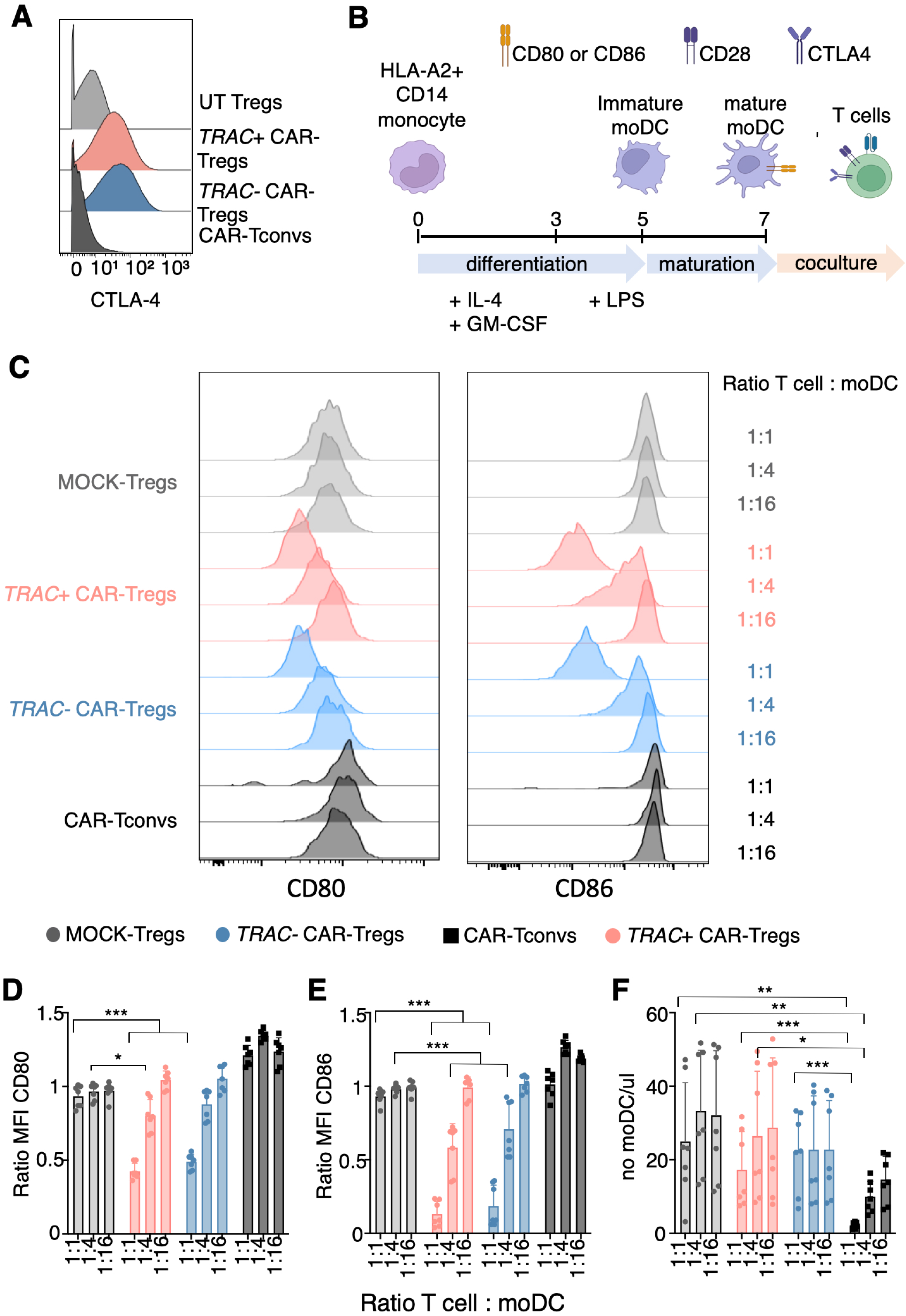
*TRAC*-deficient CAR-Tregs are as effective as TRAC-sufficient CAR-Tregs in trans-endocytosing CD80 and CD86 from the surface of moDCs. **A.C-F.** On day 16 of culture, MOCK Tregs, CAR-Tregs (either *TRAC-/-* or *TRAC+/+*) and CAR-Tconvs were put in co-culture with mature monocyte derived dendritic cells (moDC) during 40 hours with at the following T cell:moDC ratios: 1:1, 1:4, 1:16. A. Representative histograms of intracellular CTLA-4 protein expression from merged samples of n=3 different donors on day 15 of culture. **B.** Schematic design of mature moDC generation for trans-endocytosis assessment. CD14+ cells were enriched from healthy donor peripheral blood and differentiated into immature moDC in presence of IL-4 and GM-CSF before being differentiated intro mature moDC with LPS. **C-F.** Data are from n=3 donors across 3 independent experiments; bars represent the mean ± SD. **C.** Representative overlay histograms representing CD80 (left) and CD86 (right) expression at the surface of moDC for each co-culture: with MOCK Tregs (grey), *TRAC*+/+ CAR-Tregs (red), *TRAC*-/- CAR-Tregs (blue), and CAR-Tconvs (black) at the different T cell:moDC ratio. **D-E**. MFI ratios (co-cultured moDCs / moDCs alone) for CD80 (D) and CD86 (E) expression are shown. Two-tailed Mann–Whitney U tests were used to compare co-cultures with CAR-Tregs (either *TRAC*-/- or *TRAC*+/+) to MOCK-Tregs and CAR-Tconvs. At a 1:1 ratio, both CAR-Treg groups showed significant trans-endocytosis of CD80 and CD86 (p = 0.0006). At a 1:4 ratio, both CAR-Treg groups showed increased CD86 trans-endocytosis (p = 0.0006), whereas only *TRAC*+/+ CAR-Tregs exhibited increased CD80 trans-endocytosis (p = 0.011). **F.** Number of live moDCs after 40 hours of co-culture with MOCK Tregs, CAR-Tregs (*TRAC*-/- or *TRAC*+/+), or CAR-Tconvs. At a 1:1 ratio, co-culture with CAR-Tconvs led to a significant reduction in the number of live moDCs compared to MOCK Tregs (p = 0.012) and compared to both CAR-Treg groups (p = 0.0006).

To determine whether this suppressive mechanism is retained in CAR-Tregs in the absence of endogenous TCR signaling, we assessed the capacity of *TRAC*-deficient and -sufficient CAR-Tregs to mediate CTLA-4-dependent transendocytosis. At a 1:1 T cell to HLA-A2-expressing monocyte-derived Dendritic Cells (moDC) ratio, both CAR-Treg populations significantly reduced surface expression of CD80 (ratio on moDC alone: 0.42±0.05 for *TRAC*+/+ CAR-Tregs and 0.49±0.05 for *TRAC*-/- CAR-Tregs) and CD86 (0.13±0.09 for *TRAC*+/+ CAR-Tregs and 0.19±0.14 for *TRAC-/*- CAR-Tregs) on moDCs, in contrast to MOCK-Tregs (0.93±0.07 for CD80 and 0.93±0.05 for CD86) and CAR-Tconvs (1.2±0.07 for CD80 and 1.0±0.08 for CD86). This suppressive effect on CD86 expression remained statistically significant even at a 1:4 ratio (0.58±0.16 for *TRAC*+/+ CAR-Tregs and 0.7±0.19 for *TRAC*-/- CAR-Tregs). Notably, this contact-dependent suppressive function was preserved in *TRAC*-deficient CAR-Tregs, indicating that CAR signaling alone is sufficient to endow CAR-Tregs with the capacity to modulate dendritic cell immune function (**figures 4C-E and supplementary figure 5**.

Importantly, while CAR-Tconvs reduced the number of moDCs (at a 1:1 ratio: 2.2±0.84 moDC/µL), likely due to cytotoxic activity, CAR-Tregs did not exhibit this effect (17.3±10.36 moDC/µL and 22.3±10.75 moDC/µL respectively with *TRAC*+/+ and *TRAC*-/- CAR-Tregs) (**Figure 4F**). Collectively, these data demonstrate that the absence of the endogenous TCR does not impair a key mechanism of contact-dependent immune suppression mediated by CTLA-4.

### Non mitogenic CD3 monoclonal antibody selectively targets TRAC+/+ conventional and regulatory T cells

Having confirmed the absence of CD3 expression in *TRAC*-/- CAR-Tregs (**figure 1B**), we next investigated whether this change was sufficient to confer resistance to CD3 Ab-induced lymphodepletion. To this end, a 1:1 mixture of HLA-A2-targeted *TRAC*-/- GFP+ CAR-Tconvs and *TRAC*+/+ mCherry+ Luc+ CAR-Tconvs was infused into preconditioned NSG mice one day after the administration of HLA A2+ peripheral blood mononuclear cells (PBMCs) (**figure 5A**). The animals were divided into two groups: one receiving anti-CD3 antibody treatment and the other serving as untreated controls. At day 3 post-infusion, comparable numbers of splenic *TRAC*+/+ mCherry+ Luc+ CAR-Tconvs were observed in both groups (8.3x10^6^ ±4.0x10^6^ vs 8.3x10^6^ ±4.7x10^6^, p=ns). However, following CD3 Ab therapy, a significant reduction in *TRAC*+/+ CAR-Tconv numbers was observed compared to controls at day 6 (5.9x10^6^±4.4x10^6^ vs 5.8 x10^7^±3.0 x10^7^, p=0.01) and became even more pronounced by day 12 (1.2x10^6^±1.3x10^6^ vs 3,9x10^7^±1.9x10^7^, p=0.03) (**figure 5B**). We also investigated whether *TRAC*-/- GFP+ CAR-Tconvs were selectively spared and consequently enriched in the peripheral blood following CD3 Ab therapy. Consistent with this hypothesis, the TRAC-/TRAC+ cell ratio remained stable at approximately 50% in control mice but increased significantly to over 99% by day 13 in treated animals (39.7±20.8 vs 99.0±1.0, p=0.03) (**figure 5C**). Similar patterns were observed in the bone marrow and spleen (**supplementary figure 6**). These data suggest that disruption of the TCR/CD3 complex allow the selective protection from CD3 Ab lymphodepletion.

**Figure 5:**
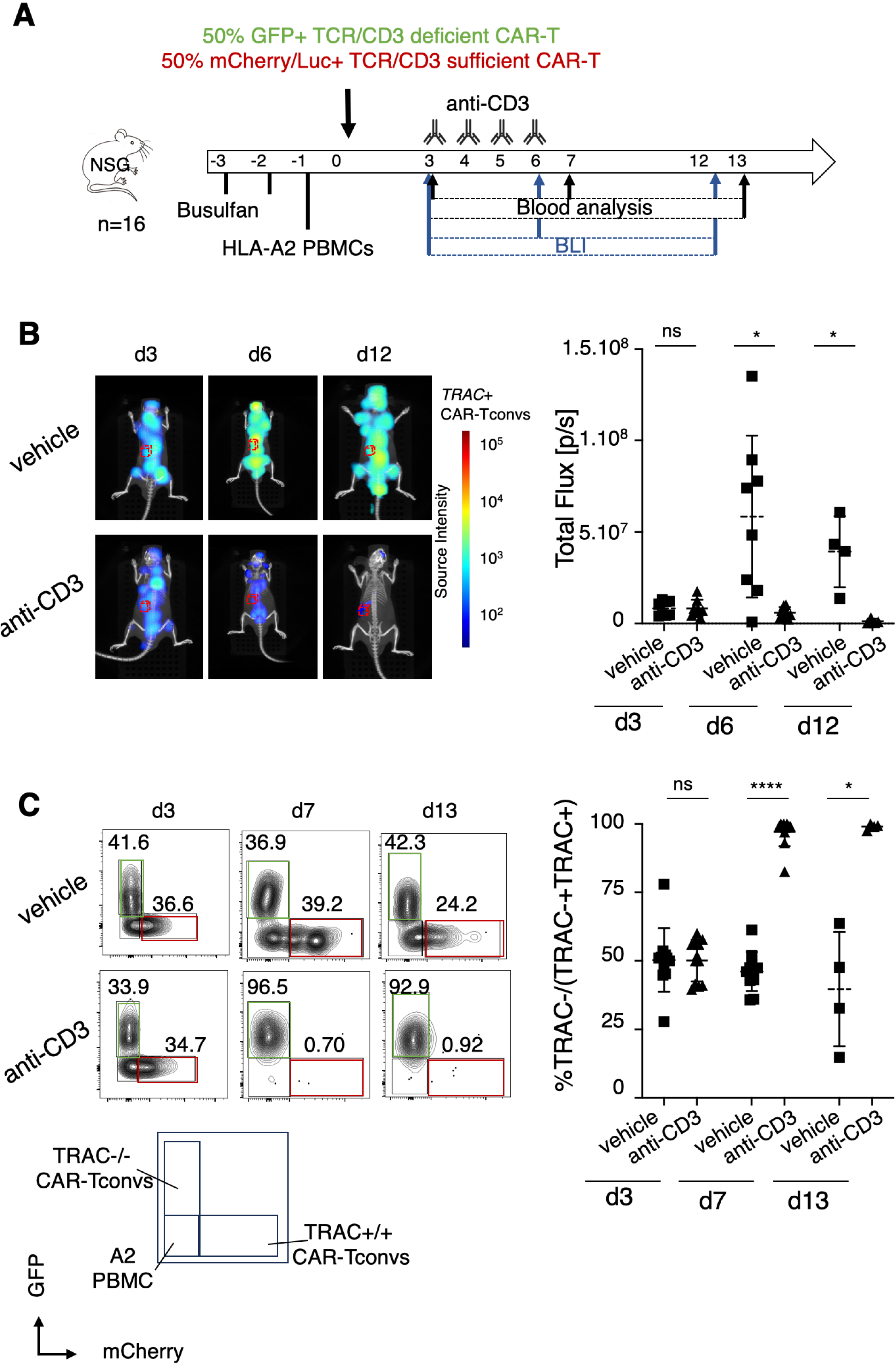
Non mitogenic anti-CD3 monoclonal antibody selectively targets *TRAC*+/+ conventional T cells. 8- to 12-week-old male NSG mice were conditioned with busulfan on days −2 and −1 before IV injection of 5x10^6^ HLA-A2+ PBMCs followed by retroorbital injection of 5×10^6^ total CAR-Tconvs (1:1 mixture of HLA-A2-targeted *TRAC*-/- GFP+ CAR-Tconvs and *TRAC*+/+ mCherry+ Luc+ CAR-Tconvs). Mice received 4 doses of 5 µg anti-CD3 on days 3, 4, 5 and 6 or were left untreated. **A.** Schematic representation of the experimental design. Mice were bled on days 3 and 7 (n=11 per group), and on day 13 (n=4 per group) for flow-cytometry analysis. Persistence of *TRAC*+/+ mCherry+ Luc+ CAR-Tconvs was assessed by bioluminescence imaging (BLI) on days 3, 6 (n=8), and 13 (N=4). **B.** 3D bioluminescence imaging tomography (DLIT) of recipient mice was performed as described in the Methods. A region of interest (ROI) corresponding to the spleen was defined, and total flux (photons/second) was quantified (right panel). Two-tailed Mann–Whitney test revealed a significant reduction in signals in anti-CD3-treated mice on day 6 (p = 0.01) and on day 12 (p = 0.029). Data are depicted as mean ± SD. **C.** Representative dot plots displaying GFP+ and mCherry+ cells gated on human CD45+ cells (left panel) and quantification of the proportion of *TRAC*-/- cells among total CAR-Tconvs cells in peripheral blood. Two-tailed Mann– Whitney test showed significant increase in *TRAC*-/- proportion in anti-CD3-treated animals on day 7 (p < 0.0001) and on day 13 (p = 0.028). Data from 2 independent experiments are depicted as mean ± SD.

We next extended this finding by demonstrating that CAR-Tregs could similarly be protected from CD3 Ab-induced depletion. To this end, 1x10^6^ CAR-Tregs, either *TRAC*+/+ or *TRAC*-/-, were transferred into preconditioned NSG mice one day after HLA A2+ PBMCs (**figure 6A**). CAR-Treg proportions in blood (**supplementary figure 7**) and their presence in the spleen (**figure 6B**) were comparable across groups prior to CD3 Ab administration (day 5). Consistent with previous results, both bioluminescence (BLI) tracking (**figure 6B**) and blood chimerism analysis (**figure 6C and supplementary figure 7**) revealed that *TRAC*-/- CAR-Tregs were unaffected by anti-CD3 administration. Comparable numbers of *TRAC*-/- CAR-Tregs in the spleen (3.54x10^5^±5.9x10^5^ vs 2.45x10^5^±2.8x10^5^) and proportion in blood (75.7±6.1% vs 67.4±12.8%) were detected respectively in the presence and absence of the antibody. In contrast, *TRAC*+/+ CAR-Tregs were effectively depleted from both lymphoid tissues and peripheral blood following CD3 Ab treatment (**figures 6B-C and supplementary figure 7**). Taken together, these findings support the potential of the combination strategy, demonstrating selective CD3 Ab-induced depletion of *TRAC*-sufficient T cells while sparing *TRAC*-deficient CAR-Tregs.

**Figure 6:**
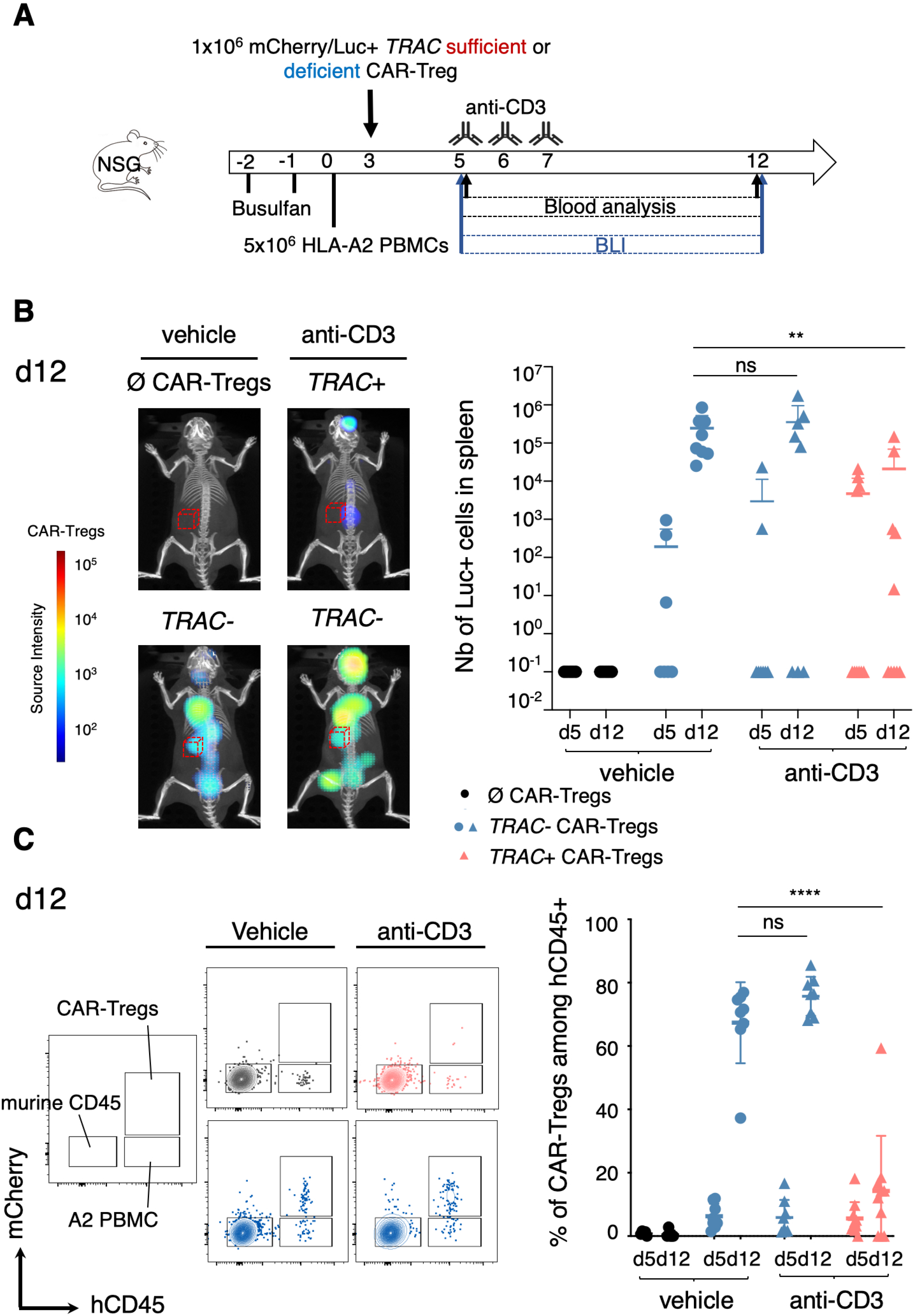
Non mitogenic anti-CD3 monoclonal antibody selectively targets *TRAC*+/+ regulatory T cells but spares their *TRAC*-/- counterpart. 8- to 12-week-old male NSG mice were conditioned with busulfan on days −2 and −1 before IV injection of 5x10^6^ HLA-A2+ PBMCs followed by retroorbital injection of 1×10^6^ Luc+ mCherry+ CAR-Tregs, either *TRAC*+/+ or *TRAC*-/-. Mice received 3 doses of 3 µg anti-CD3 on days 5, 6 and 7 or were left untreated (at least n=8 mice per group). **A.** Schematic representation of the experimental design. Mice were bled for flow-cytometry analysis and persistence of *TRAC*+/+ or *TRAC*-/- mCherry+ Luc+ CAR-Tregs was assessed by BLI on days 5 and 12. **B-C.** Day 12 data from 2 independent experiments are shown as mean ± SD. **B.** Representative 3D images (left panel) and total number of Luc+ cells was quantified by DLIT acquisition (right panel). A two-tailed Mann– Whitney test revealed a significant reduction in the number of *TRAC*+/+ CAR-Tregs of anti-CD3-treated mice (p = 0.019) while no significant difference was observed for *TRAC*-/- CAR- Tregs (p = 0.72). **C.** Representative dot plots showing mCherry+ and human CD45+ cells (left panel) and quantification of the proportion of CAR-Tregs (*TRAC*+/+ or *TRAC*-/-) among human CD45 cells in peripheral blood (right panel). A two-tailed Mann–Whitney test showed significant decrease in the proportion of *TRAC*+/+ CAR-Tregs in anti-CD3-treated animals compared to *TRAC*-/- CAR-Tregs (p < 0.0001).

### CD3 monoclonal antibody therapy penetrates peripheral tissues and depletes alloreactive T cells in situ

We previously demonstrated the selective depletion of *TRAC*+/+ CAR-Tconvs in blood and lymphoid organs. We next investigated whether CD3 Ab treatment could also target alloreactive T cells within a graft. Given the critical role of tissue-infiltrating Tconvs in local immune responses and graft rejection, we assessed the ability of anti-CD3 to access target organs and deplete alloreactive T cells *in situ*, while sparing *TRAC*-/- CAR-Tregs.

To this end, we established an *in vivo* dual-luciferase imaging system enabling simultaneous tracking of CBG+ *TRAC+*/+ CAR-Tconvs and Fluc+ *TRAC-*/- CAR-Tregs. As expected, CBG+ cells emitted at 540 nm, while Fluc+ exhibited a distinct peak at 620nm, both *in vitro* and *in vivo* (**supplementary figures 1 and 8A-C)**. Spectral unmixing using the IVIS Spectrum system allowed robust separation of the two luciferase signals and independent quantification of each population. Signal intensities correlated with injected cell ratios, confirming the reliability of the dual-luciferase system (**supplementary figure 8C**).

To evaluate CD3 Ab mediated depletion in tissues, *TRAC+*/+ CAR-Tconvs and *TRAC*-/- CAR-Tregs were co-injected into mice bearing HLA-A2 allogenic skin grafts (**figure 7A)**. By day 3, dual-luciferase imaging revealed rapid homing of both cell populations to the graft site, as indicated by detectable CBG (*TRAC*+/+ CAR-Tconvs) and Fluc (*TRAC*-/- CAR-Tregs) signals (**figure 7B, left panel**). Following anti-CD3 administration, the CBG signal markedly decreased, reflecting depletion of *TRAC*+/+ CAR-Tconvs. This resulted in a significant increase in the Fluc/CBG signal ratio (1.98±1.91 in controls vs 12.9±12.7 in treated mice), indicating enrichment of *TRAC*-/- CAR-Tregs at the graft site (**figure 7B, right panel**). In contrast, untreated mice maintained a stable signal ratio, consistent with persistence of both cell populations. These findings demonstrate that non-mitogenic anti-CD3 therapy can penetrate tissues and selectively deplete *TRAC*+/+ alloreactive Tconvs, while leaving *TRAC*-/- CAR- Tregs unaffected.

**Figure 7:**
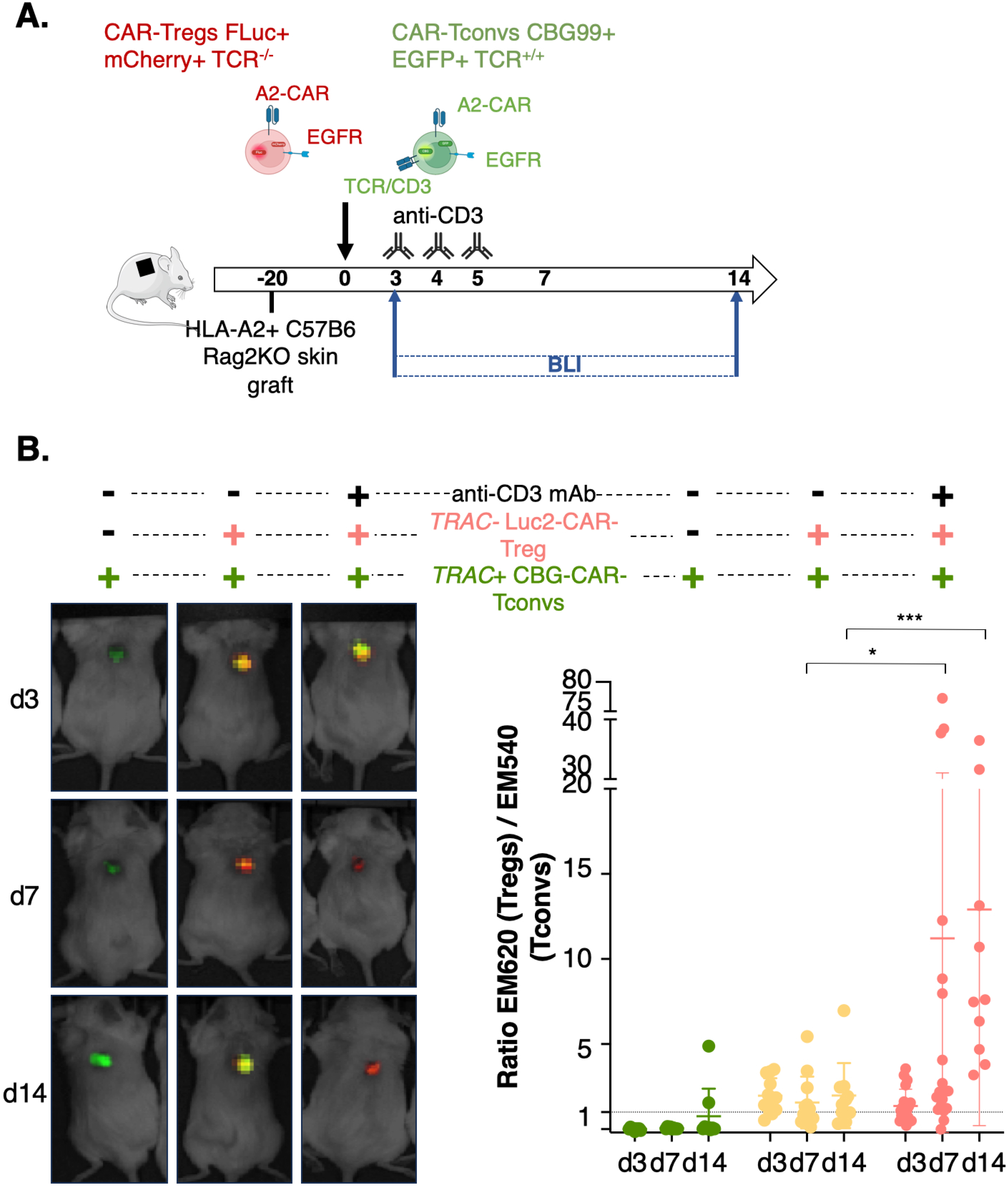
Anti-CD3 therapy penetrates peripheral tissues and depletes alloreactive T cells in situ while preserving *TRAC*-/- CAR-Tregs. **A.** Schematic representation of the experimental design. **B.** Mice grafted with HLA-A2+ allogeneic skin received 2x10^6^ *TRAC*-/- Fluc+ CAR-Tregs and 2x10^6^ *TRAC+/+* CBG+ CAR-Tconvs 3 weeks post-surgery. On day 3 days post-injection, mice received 5µg anti-CD3 (n=19) or were left untreated (n=12). (Left panel) Representative 2D composite images on days 3 (pre-treatment), 7 and 14. (Right panel) Quantification of Fluc+/CBG+ signal ratio representing the relative proportions of TRAC-/- CAR-Tregs versus TRAC+/+ CAR-Tconvs over time. A two-tailed Mann–Whitney test showed significant enrichment of *TRAC*-/- CAR-Tregs in anti-CD3-treated animals compared to untreated animals on day 7 (p = 0.0317) and day 14 (p = 0.0001). Data from 4 independent experiments are depicted as mean ± SD.

### Non-mitogenic CD3 monoclonal antibodies selectively induce exhaustion and apoptosis in TRAC+/+ conventional T cells, while preserving the transcriptomic profile of TRAC-/- CAR-Tregs

The non-mitogenic Fc mutated CD3 Ab otelixizumab was engineered (humanized and Fc-mutated) from the original rat YTH 12.5 monoclonal antibody to reduce the risk of cytokine release syndrome and immunization^24,25^. Although this antibody induces substantially lower cytokine production, both *in vivo* and *in vitro* studies have reported transient elevations in pro-inflammatory cytokines, including IFNγ, TNFα, interleukin (IL-)17 and IL-6. To assess potential bystander effects of CD3 Ab-induced cytokine release on CAR-Treg biology *in vivo*, we performed scRNA sequencing on *TRAC-*/- CAR-Tregs and *TRAC+*/+ CAR-Tconvs isolated from spleens of mice treated or untreated with non-mitogenic anti-CD3 (**figure 8A**).

**Figure 8:**
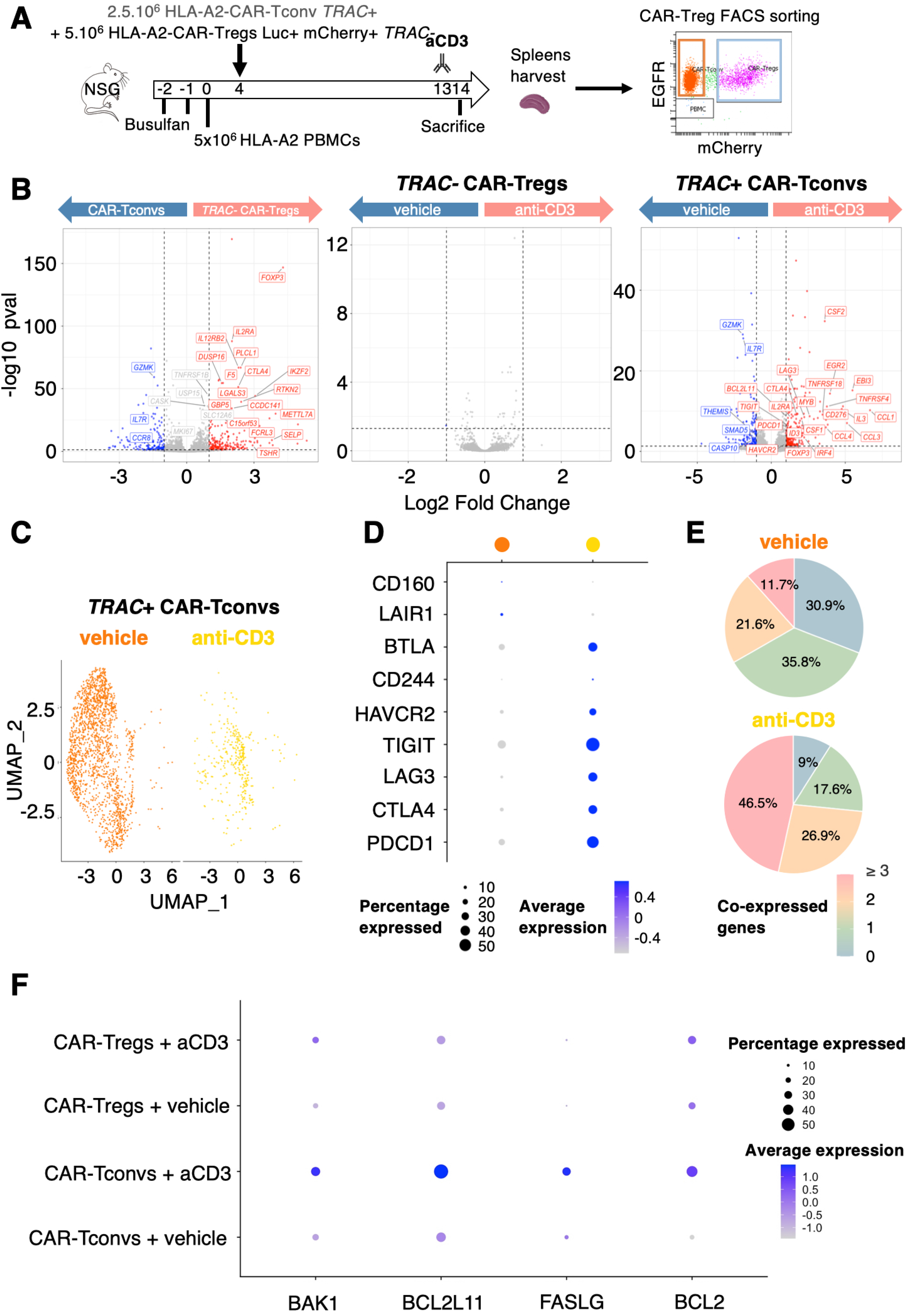
A non-mitogenic anti-CD3 treatment selectively induces exhaustion and apoptosis in *TRAC*+/+ conventional T cells, without compromising the transcriptomic integrity of *TRAC-*/- CAR-Tregs. *TRAC*+/+ CAR-Tconvs and *TRAC*-/- CAR-Tregs were collected 12 hours after anti-CD3 administration from the spleens of 8- to 12- week-old preconditioned NSG male mice injected with HLA-A2+ PBMC and a 1:2 ratio of *TRAC*+/+ CAR-Tconvs to *TRAC*-/- CAR-Tregs. **A.** Schematic representation of the experimental design. **B-F.** Single-cell RNA-seq analysis from one experiment with 3 mice per group **B.** Volcano plots generated from a pseudobulk analysis comparing (left) *TRAC-* CAR-Tregs with *TRAC+* CAR-Tconvs from untreated mice, (middle) *TRAC-* CAR-Tregs from anti-CD3 treated with untreated mice and (right) *TRAC+* CAR-Tconvs from anti-CD3 treated with untreated mice. **C.** UMAP showing distribution of cells: *TRAC*+/+ CAR-Tconvs from anti-CD3-treated mice (yellow) and *TRAC*+/+ CAR-Tconvs from untreated mice (orange) clustered differently. **D.** Dot plots generated with *Seurat* package showing the expression of selected exhaustion-related genes and **E.** pie charts depicting the proportion of cells expressing 0, 1 or co-expressing 2 or ≥3 exhaustion genes, comparing *TRAC*+/+ CAR-Tconvs from anti-CD3-treated vs untreated mice. **F**. Dot plots generated with *Seurat* package showing the expression of selected apoptosis-related genes.

*TRAC*-/- CAR-Tregs isolated from untreated mice exhibited a canonical Treg-core signature characterized by expression of *FOXP3, IKZF2, IL2RA* and *CTLA-4*, when compared with CAR-Tconvs. Importantly, comparative analysis of *TRAC-/-* CAR-Tregs from CD3 Ab treated versus untreated mice showed no significant differentially expressed genes, indicating that the absence of TCR confers resistance to treatment-induced transcriptional alterations (**figure 8B**).

In contrast, similar analysis of *TRAC*+/+ CAR-Tconvs revealed multiple differentially expressed genes, reflecting a robust transcriptional response to CD3 Ab that drove treatment-dependent cell clustering (**figures 8B and 8C**). Notably, *TRAC*+/+ CAR-Tconvs from CD3 Ab treated mice showed upregulation of genes associated with T cell exhaustion (*PDCD1, TIGIT, LAG3, HAVCR2, LAIR2, CD200*, and *CD276*) as well as genes related to regulatory functions (*FOXP3, IL2RA, CTLA4, ENTPD1, EBI3, FURIN, TNFRSF18, TNFRSF4, TNFRSF9*) (**figure 8D**). Quantitative analysis revealed that the proportion of CAR-Tconvs expressing no exhaustion markers decreased from 30.9% to 9.0% following CD3 Ab treatment, while the proportion of those expressing three or more increased fourfold (**figure 8E**). In contrast, exhaustion markers expression remained stable in *TRAC*-/- CAR-Tregs (**supplementary figures 9A-B**). In line with CD3 Ab-induced T cell depletion (**figures 5-7**), CAR-Tconv cells showed increased expression of genes involved in both extrinsic (FASLG) and intrinsic (BAK1, BCL2L11) apoptotic pathways following CD3 therapy (**figure 8F**).

Collectively, these findings demonstrate that non-mitogenic anti-CD3 selectively induces exhaustion and apoptosis in *TRAC*+/+ conventional T cells, while preserving the transcriptional integrity of *TRAC*-/- CAR-Tregs.

## DISCUSSION

Our study highlights the significant therapeutic potential of combining non-mitogenic CD3 Ab with *TRAC*-/- CAR-Tregs. Various strategies aimed at promoting immune tolerance through the enhancement of Treg and inhibition of effector T cells have shown efficacy in experimental transplantation models, particularly in rodents. These include the classical combination of CD40-CD40L and CD28-B7 blockade^26^, as well as other synergistic regimens such as IL-2 agonist, IL-15 antagonist, and mTOR inhibition^27^. The use of anti-CD45RC antibodies follows a similar rationale by targeting the CD45RC^high^ Tconvs while sparing CD45RC^low^ Tregs^28^.

However, such strategies have predominantly been effective in immunologically naïve recipients, lacking pre-existing activated/memory alloreactive T cells. In contrast, human transplant recipients typically possess pre-formed alloreactive memory T cells, often arising from heterologous immune responses. These cells are characteristically resistant to costimulatory blockade.

This key immunological distinction likely underpins the limited clinical success of many tolerance-inducing immunomodulatory protocols10,29,30. Consequently, the most effective tolerogenic strategies in clinical practice have incorporated profound lymphodepletion, achieved through agents such as anti-CD2 antibody, total lymphoid irradiation or post-transplant cyclophosphamide^7,14^. Our strategy preferentially targets activated T cells, which are particularly enriched in alloreactive populations during the immediate post-transplant period, while preserving a stealth, donor-targeted regulatory cell therapy product. This *in vivo* advantage-based approach shifts the immune balance toward regulation and may substantially enhance the efficacy of CAR-Treg in clinical settings. Moreover, it has the potential to reduce the number of regulatory cells required for therapeutic efficacy, thereby lowering associated manufacturing costs.

CD3 Abs exhibit the unique ability to restore or induce immune tolerance even after the priming of a deleterious immune response. This characteristic implies efficacy against activated effector and memory T cells. In patients with newly diagnosed type 1 diabetes, a 6-day-course of otelixizumab conferred a durable benefit on insulin requirement lasting over four years^31,32^. Such a prolonged effect cannot be solely attributed to transient immunosuppression but rather suggests the induction of genuine tolerogenic mechanisms. The efficacy of another non-mitogenic CD3 Ab, teplizumab, in preventing the progression from stage 2 to stage 3 type 1 diabetes has been well established^33^. Preclinical models have shed light on the mechanisms of action of these second-generation CD3 Abs. They selectively induce apoptosis in activated T lymphocytes while sparing resting T cells, thus preserving a naïve repertoire capable of mounting effective anti-infectious responses^16,34,35^. Notably, these antibodies do not induce sustained lymphopenia^35^, in contrast to anti-lymphocyte sera or alemtuzumab (CD52 monoclonal antibody). Instead, transient margination of lymphocytes, particularly within the splanchnic circulation during treatment, has been observed and is rapidly reversible upon cessation^36^.

Remarkably, effector T cells that survive CD3 Ab treatment acquire a dysfunctional phenotype suggestive of exhaustion/anergy. Our findings in a humanized mouse model corroborate and extend those observed in diabetic patients^37,38^ and in murine models^39^. The induction of the concomitant expression of several co-inhibitory molecules by CAR-Tconvs after CD3 Ab therapy is indicative of an exhausted phenotype, similar to what is observed in type 1 diabetic patients^37,38^.

The tolerogenic properties of CD3 Abs extend beyond the depletion and acquired dysfunction of T cells. The phagocytosis of T-cell-derived apoptotic bodies by macrophages in lymphoid tissues draining the target organs of pathogenic lymphocytes promotes TGF-β secretion^17^, thereby fostering a local microenvironment conductive to the induction of Tregs^18^.

Although Tregs show partial resistance to CD3 Ab-mediated depletion compared to Tconvs^40^ *ex vivo* production of CAR-Tregs requires activation, which sensitizes CD3/TCR-expressing cells to pro-apoptotic effects. To address this, genetic disruption of the CD3/TCR complex can prevent the unintended loss of a portion of the cell therapy product following CD3 Ab administration.

A key determinant of the tolerogenic effect of CD3 Ab therapy lies in the timing of its administration relative to the activation of pathogenic T cells. In preclinical transplantation models (pancreatic islets, heart), it has been well established that donor-specific tolerance require administration of CD3 Abs after the activation of alloreactive T cells, typically 3-7 days post-transplantation^16,34^. This finding is consistent with observations in autoimmune disease models (type 1 diabetes, Experimental autoimmune encephalomyelitis), where CD3 Ab treatment durably restores tolerance only when administered after the onset of initial symptoms. When given earlier, it merely delays disease progression without achieving lasting tolerance^41,42^. Our experimental designs were accordingly adapted, relying on a prior *in vivo* activation of HLA-A2-targeted CAR-Tconv cells by HLA-A2+ cells before the administration of otelixizumab.

In clinical transplantation, however, identifying the optimal therapeutic window of CD3 Ab remains challenging, particularly due to the presence of memory alloreactive T cells, which can become reactivated more rapidly than naïve T cells in SPF-housed murine models. A recent study suggests that in sensitized murine recipients, CD3 Ab therapy must be initiated earlier to achieve efficacy^43^. These findings support the notion that, in clinical settings, initiating treatment on day 2 rather than day 5 post-transplantation may be more appropriate. Importantly, in pancreatic islet transplantation, teplizumab offers the dual benefit of targeting both autoimmune and alloimmune T cell responses, with promising clinical potential.

We sought to investigate the impact of TCR deletion on the transcriptomic, epigenetic, phenotypic and functional properties of Tregs. The homeostatic maintenance of Tregs is known to rely on interactions between the TCR and MHC class II molecules^44^. Inducible ablation of the TCR in Tregs leads to a disruption of approximately 25% of their transcriptomic signature and impairs their suppressive function^45^. Notably, Tregs engage their TCR more strongly than Tconv, even when sharing the same TCR, suggesting an intrinsic ability to compete for antigen recognition and thereby sustain their regulatory identity^46^. These findings have raised concerns that deletion of the TCR/CD3 complex in Tregs might negatively impact their biology and function. Nevertheless, our data demonstrate that *TRAC*-/- CAR-Tregs preserve both their regulatory identity and functional competence after 16 days in culture. This suggests that the CAR delivers a signal through its intracellular CD3ζ and/or CD28 domains, sufficient to maintain the minimal activation threshold required for preserving the regulatory program. Our group and others have previously shown that CAR constructs incorporating a CD28 co-stimulatory domain are superior to those with alternative domains, such as 4-1BB, for CAR-Treg engineering^4,5^. Indeed, CD28 signaling is essential for Treg homeostasis^47^ and for maintaining their epigenetic landscape^48^.

The combined strategy developed in this study offers assurances for successful clinical translation. It integrates several therapies and technologies previously tested in humans, each demonstrating a favorable efficacy-to-safety ratio. Targeted gene editing using nucleases, such as TALENs (transcription activator-like effector nucleases) or CRISPR-Cas9, has already been implemented clinically to engineer CAR-T cells. TCR deletion reduces the risk of off-target effects and eliminate the potential for GVHD in the context of allogeneic CAR-T cells^49,50^. *CD52* gene knockout confers resistance to lymphodepletion by alemtuzumab and provides a selective *in vivo* survival advantage to the modified cells^49,51^. Several CAR-Treg trials are currently underway, including two in solid organ transplantation (kidney and liver), employing autologous CAR-Tregs targeting donor HLA-A2 antigens (STEADFAST, NCT04817774; LIBERATE, NCT05234190).

Our study has several limitations with regard to clinical translation. The adoptive transfer of human PBMCs into immunodeficient murine hosts does not recapitulate the full complexity of the human immune system, particularly in terms of cellular diversity and the spatiotemporal orchestration of immune responses within lymphoid organs. For example, our model does not capture the phagocytosis of apoptotic lymphocytes by macrophages, a process that contributes to a local anti-inflammatory environment dependent on TGF-β^17^. Moreover, the transfer of mature, non-renewable T cells in the absence of thymic output, precludes long-term assessment of tolerance mechanisms and effective prevention of chronic rejection. Indeed, the depletion of pathogenic T cells by CD3 Ab eliminates the source of IL-2 necessary for sustained maintenance of functional Tregs. Consequently, we were unable to adequately assess the long-term persistence, stability, and functionality of human CAR-Tregs in NSG host following CD3 Ab treatment. In this context, rendering CAR-Tregs autonomous in IL-2 signaling may circumvent the risk of destabilization in IL-2-deprived environments^52,53^. A recent study demonstrated enhanced proliferation, survival, and function of Tregs engineered to express an IL-2 mutein^54^.

In summary, our combinatorial strategy confers a selective *in vivo* advantage to donor-antigen-specific CAR-Tregs over alloreactive Tconvs, thereby shifting the immunological balance toward tolerance. This approach may enhance the capacity of CAR-Tregs to induce robust and infectious tolerance^55^, potentially sustaining immune regulation even after the infused CAR-Tregs have been cleared^56^.

## METHODS

### Construct design

The selected scFv for the CAR construct was fused to a stalk region from the human CD8α hinge, the transmembrane domain of human CD8, the CD28 CSD, and the signaling domain of human CD3ζ. As described previously^5^, EGFRt was cloned downstream of a T2A self-cleavage peptide in the second gene position to serve as a reporter. A similar construction without the extracellular domain is used as a MOCK condition (supplementary figure 1A). The lentiviral plasmid encoding firefly luciferase (Fluc) protein (pEFS-Fluc-mCherry) was kindly provided by Dr. Matthias Titeux. The click beetle green protein (CBG) LV (pEFS-CBG-GFP) was derived from the Fluc vector.

### Cell source and purification of human Tregs

Naïve Tregs and naive Tconvs were obtained from cytapheresis products collected from healthy HLA-A2/A28-negative donors under 40 years of age, as previously described^5^. CD4+ T cells were enriched using the EasySep CD4+Enrichment kit (Stem Cell, Canada). Naïve Tregs and Tconvs were respectively sorted on a FACS Aria II (BD Biosciences, New York), according to the phenotypes CD4^+^CD25^Hi^CD127^−^CD45RA^+^CD45RO^−^ and CD4^+^CD45RA^+^CD25^−^, achieving a purity exceeding 98%. Sorted cells were cultured in X-VIVO® 20 medium supplemented with 10% human AB serum (Biowest, France) and 1000 UI/mL human IL-2 (Proleukin, Novartis, Switzerland). The following day, cells were stimulated with T cell TransAct (Miltenyi, Germany) at a 1:50 dilution.

### T-cell transduction and expansion

One day after activation, Tregs and Tconvs were transduced with the CAR-encoding LV at a multiplicity of infection (MOI) of 30 viral particles per cell, in combination with either the Fluc-mCherry-encoding LV (MOI 5) or CBG-GFP-encoding LV (MOI 40) as previously described^5^. On day 5 post transduction, CD4+ EGFRt+ cells were sorted using a FACSAria^TM^ II and subsequently restimulated at a 1:1 ratio with B cells enriched from HLA-A2+, CD3-depleted splenocytes. UT T cells were restimulated with T cell TransAct^TM^ (Miltenyi, Germany) at a 1:100 dilution. T cells were then cultured and expanded in complete medium supplemented with IL-2 (1000 UI/mL) for 7 days, prior to phenotypic, transcriptomic, epigenetic, and functional analysis. This protocol was deliberately shortened to minimize Treg destabilization associated with prolonged *in vitro* culture and to reduce the future costs inherent to extended GMP-grade manufacturing.

HLA-typed, CD3-depleted splenocytes were isolated from spleens obtained from deceased organ donors, in collaboration with the Regional Histocompatibility Laboratory and National Biomedicine Agency. B cells were then enriched using an EasySep negative selection kit (Stem Cell, Canada).

### TRAC deletion using CRISPR/Cas9 genome editing and nucleofection

Guide RNA (gRNA) target sites within the human *TRAC* exon were identified as described in^5^. The *TRAC*-targeting and non-targeting CRISPR RNA (crRNA) were purchased from Thermofisher, Waltham and sequences are displayed in **supplementary table 3**. Three days post-activation, cells were nucleofected^5^.

### Phenotypic characterization of gene-edited Tregs

Surface staining was performed using monoclonal antibodies listed in supplemental Table 4. For intracellular staining, cells were first stained with a fixable viability dye (FVD, Thermo Fisher Scientific, Waltham), followed by surface staining. Cells were then fixed, permeabilized, and stained using the FOXP3/transcription factor staining buffer set (eBioscience, San Diego), according to the manufacturer’s instructions. Samples were analyzed using a BD LSRFortessa^TM^ X-20 analyzer (BD Biosciences, New York) and data were processed with FlowJo 10.10 (BD Biosciences, New York).

For high-dimensional phenotypic profiling by mass cytometry, 1x10^6^ cells were stained with Cell-ID™ Cisplatin (Fluidigm, San Fransisco) and samples from each donor were individually barcoded to enable multiplexed acquisition. Samples were pooled for surface and intracellular staining according to the workflow presented in the **supplementary figure 10** and **supplementary table 2**. Antibodies were either obtained pre-conjugated, or manually conjugated using the Maxpar X8 Antibody Labelling Kit (Fluidigm, San Fransisco). Samples (mixed with 10% EQ^TM^ Four Element Calibration Beads for normalization) were acquired on the Helios^TM^ mass cytometer (Fluidigm, San Fransisco) equipped with CyTOF® software version 6.7.1014 by the Cytometry Pitié Salpêtrière (CyPS) Platform. Mass cytometry data were analyzed using the OMIQ data analysis software (www.omiq.ai).

### Activation assay

UT Treg, scramble controls and *TRAC*-edited CAR-Tregs were seeded into 96-well flat-bottom plates at 1.10^5^ cells per well and cocultured with either HLA-A2+ or HLA-A2-B cells (enriched from splenocytes) at a 1:1 ratio. Expression of activation markers (listed in **supplementary table 4**) was assessed by flow cytometry after 40 hours of co-culture.

### Bulk RNA sequencing sample preparation

Scramble- or *TRAC*-edited CAR-Tregs and CAR-Tconvs were sorted at day 15 of culture on the viable EGFR-expressing cells (supplementary figure 1C) and stored at -80°C in RLT buffer (Qiagen, Germany). Total RNA was extracted using RNeasy Mini Kit (Qiagen, Germany) according to the manufacturer’s instructions. RNA quality and integrity were assessed using a Fragment Analyzer 5200 (Agilent Technologies, Santa Clara), and samples with an RNA integrity number RQN >8 were selected for sequencing. Bulk RNA sequencing was performed at the SFR Necker genomics platform using the Illumina NovaSeq 6000 system. FASTQ files were generated by demultiplexing and aligned to the human reference genome (Ensembl GRCh38) using HISAT2. All samples passed the quality control, assessed using Picard tools. For the downstream analyses, genes with average read counts ≥20 in ≥20% of samples were retained. Differential expression analysis was performed using the DESeq2 package in R, and genes with a p-value < 0.05 were considered significantly differentially expressed. An activation human Treg signature was extracted from a published human dataset (GSE76598)^22^ downloaded through the Gene Expression Omnnibus (GEO) (https://www.ncbi.nlm.nih.gov/geo/). Up and downregulated transcripts from the comparison “stimulated Tregs vs stimulated Tconvs” were selected.

### Demethylation analysis

Bisulfite conversion, pyrosequencing and subsequent analyses were performed by EpigenDX company. Frozen cell pellets (5.10^5^ cells) from UT Treg, scramble controls, *TRAC*-edited CAR-Tregs and CAR-Tconvs were analyzed using the human N70V3P15 panel. Results are reported as the percentage of T cells with a demethylated TSDR region. For female patients, this percentage was corrected by a factor of two to account for X-chromosome inactivation, which results in one of the two TSDR alleles being constitutively methylated.

### TransEndocytosis assay

HLA-A2-expressing CD14+ monocytes were enriched from human PBMCs using the EasySep™ Human Monocyte Enrichment Kit (StemCell, Canada) and cultured with 50 ng/mL IL-4 and 10 ng/mL GM-CSF for five days. Immature dendritic cells were then activated with 50 ng/ml LPS (Sigma-Aldrich, Saint-Louis) for two days. CD80/CD86 expressing moDC were seeded in co-culture with either MOCK-Tregs (transduced with an empty LV), *TRAC*+/+ CAR-Tregs, *TRAC*-/- CAR-Tregs or CAR-Tconvs on day 15 of culture in 96 well U-bottom plate at the following T cell: moDC ratios: 1:1, 1:4, 1:16. After 40 hours incubation at 37°C, viability and CD80/CD86 expression were assessed by flow cytometry (NovoCyte Penteon, Agilent) using the antibodies listed in **supplementary table 4**.

### In vivo anti-CD3-mediated depletion assay

Male NSG mice (8-12 weeks old), either bred in-house or purchased from Charles River Laboratories, France, received intraperitoneally injections of busulfan (25 mg/kg; Merck, Saint-Louis) on days −2 and −1 prior to human PBMC transfer. On day 0, mice were injected intravenously (via the tail vein) with 5.10^6^ HLA-A2+ PBMCs. At the designated time points, mice were injected via the retro-orbital venous sinus with 2-3.10^6^ CAR-Tconvs and/or 1.10^6^ CAR-Tregs, with or without *TRAC* gene editing. Mice received subsequently 3-5 µg of Otelixuzumab (anti-CD3) intravenously once daily for 3 to 4 consecutive days. At the indicated time points, peripheral blood was collected from the venous sinus under isoflurane anesthesia and tetracaine-based analgesia. Spleen and bone marrow were also harvested. Red blood cells were lysed using RBC lysis buffer (Ozyme, France), and leukocytes were analyzed by flow cytometry.

### In vivo cell tracking

On day 14 of culture, Fluc-expressing Tregs and CBG luciferase-expressing Tconvs were intravenously injected into either busulfan-conditioned NSG mice or skin-grafted recipient mice. For bioluminescence imaging, D-luciferin potassium salt (150 mg/kg, Revvity, Waltham) was administered intraperitoneally prior to imaging under isoflurane anesthesia.

### 3D imaging using IVIS Spectrum

Images were acquired using the IVIS SpectrumCT imaging system (Revvity, Waltham) with the DLIT (Diffuse Luminescent Imaging Tomography) tool and data were analyzed using Living Image® software (IVIS Imaging Systems) as previously described^5^.

### Spectral Unmixing imaging using IVIS Spectrum

Bioluminescence imaging (BLI) was performed on the IVIS® Spectrum small animal imaging system (Revvity, Waltham). Emission signals were captured using spectral unmixing with band-pass filters set at 540, 560, 620 and 640 nm. Image acquisitions occurred 15 minutes after substrate administration, with a minimum exposure time of 30 seconds per image. Binning was adjusted between 4 et 16, and f/stop ranged from 1 to 8. Data were analyzed using Living Image® software version 4.3 (Perkin Elmer), employing a custom spectral library established during a pilot study. Signal normalization was performed using control animals that did not receive bioluminescent cells, following Revvity’s recommended procedures.

### Single cell RNA sequencing (scRNA-Seq)

HLA-A2-targetted *TRAC*-deficient CAR-Tregs and *TRAC*-sufficient CAR-Tconvs were injected to preconditioned NSG mice 4 days after HLA-A2-expressing human PBMCs. A single dose of 5 µg anti-CD3 antibody was administered intravenously to half of the mice. CAR-Tregs and CAR-Tconvs were isolated from spleens 12 hours later for single cell analysis using the Rhapsody platform. Each condition was labelled using separate Sample Tag (BD Bioscience, New York) before multiplexing. Single cell capture and library preparation were performed using the microwell-based BD Rhapsody scRNA-Seq platform with the BD Rhapsody Whole Transcriptome Analysis (WTA) reagent kit (BD Biosciences, New York). RNA transcripts and sample tags libraries were prepared according to the manufacturer’s instructions. Quality controls was conducted using the Promega Quantus fluorometer and the Agilent Bioanalyzer. Libraries were sequenced on the NextSeq 2000 device (300-cycle kit) (Illumina, San Diego). The FASTQ files were processed using the BD Rhapsody WTA analysis pipeline with the GRCh38-PhiX reference human genome (Seven Bridges). Downstream analysis was performed using R (v4.4), Seurat (v5.3.0) in R studio (v2024.12.0+467). Cells with fewer than 1,000 or more than 5,000 detected features, or with >30% mitochondrial gene content, were excluded from the analysis. Pseudo-bulk analysis was performed using scRNA-seq data processed with Seurat. Differential expression analysis between anti-CD3–treated and control CAR-Tconvs was performed using DESeq2. Lowly expressed genes were filtered prior to analysis, and significance was defined by an adjusted p-value < 0.05 and an absolute log2 fold change > 1.

### Skin graft

Allogeneic tail skin 1cm^2^ biopsy from HLA-A2 Rag2KO C57BL/6 (bred in house) was surgically transplanted to 8- to 12-week-old NSG mice (bred in house or purchased from Charles River, France) in the dorsal area. A compression bandage was applied to protect the graft for 10 days and was subsequently removed along with the eight stitches securing the graft.

### Statistical analysis

Results are presented as mean ±SD. Frequencies of categorical variables are reported as absolute numbers and percentages. Statistical analyses were performed using GraphPad Prism software (version 10.4.0; GraphPad Software). For comparisons of two groups in *in vitro* experiments, the nonparametric two-tailed Mann–Whitney test was used. p-values < 0.05 were considered statistically significant. Principal component analysis (PCA) and volcano plots were generated using RStudio (version 2024.12.0+467) with the dplyr, ggrepel and ggplot2 packages.

### Ethical approvals

This study was conducted in accordance with all relevant ethical regulations. Spleen collection from deceased organ donors was approved by a local Institutional Review Board (Project no.: 2016-12-01; approved on December 5, 2016). All animal procedures were approved by the “Services Vétérinaires de la Préfecture de Police de Paris” and by the “Comité d’Ethique en matière d’Expérimentation Animale Paris Descartes (CEEA 34)” under authorization numbers APAFIS #33303-2021081017535587 v3 and APAFIS #43552-202212131845937 v10 (Université Paris Cité, Paris, France).

## DATA AVAILABILITY

The authors declare that all data supporting the findings of this study are available in the article and its Supplementary Information Files, or upon request from the corresponding author. The data underlying all the figures are provided as a source data file. The transcriptomic data reported in this study have been deposited in the ArrayExpress database under accession no. E-MTAB-16484 and n° E-MTAB-16542. Protein expression from CYTOF data have been deposited to the zenodo database with the identifier 10.5281/zenodo.17965659.

## ACKNOWLEDGEMENT

The authors are grateful to Sophie Candon, Yannick Muller, Eliane Piaggio, and Sylvaine You for their critical and helpful feedback on the manuscript. We thank Jérôme Mégret and colleagues at the Plateforme de cytométrie; O. Claude, A. Legrand, J.C. Beche and colleagues at the Laboratoire d’Expérimentation Animale et Transgénèse and C. Bole and colleague at the Plateforme de Génomique of the Structure Fédérative de Recherche Necker, Paris; G. Froment, A. Thibaut and C. Costa from the AniRA lentivectors production facility at the CELPHEDIA Infrastructure and SFR BioSciences (UMS3444/CNRS/US8/Inserm, ENS Lyon, UCBL); the Etablissement Français du Sang for blood supply from healthy donors, B. Hoareau, A. Vinit and A. Chapel from the Mass Cytometry Platform of La Pitié-Salpêtrière Hospital (CyPS), the iGenSeq platform of the Institut du Cerveau for sequencing of the scRNA-seq libraries; SIRION biotech GmbH for providing LentiBOOST^TM^. T.B. and S.C. were supported by the Fondation Emmanuel Boussard. The study was also funded by the Fondation Recherche Medicale (PME202406019177), Fondation Day-Solvay and Agence Nationale de la Recherche (DAISY, ANR-20-CE18-0004; MAINTREG ANR-21-CE17-0015; ARMSTRONG, ANR-24-CE18-6414). The IVIS spectrum used in this study was purchased with a grant from the DIMThérapie Génique/Région Ile-de-France. T.B. created graphical abstract from biorender.com.

## AUTHORS CONTRIBUTIONS

T.B., N.A., S.C. and J.Z. conceived and designed the experiments and wrote the manuscript. T.B., N.A., S.C. and A.G. performed the experiments and analyzed the data. A.H. generated the scRNA-seq libraries. J.H. and J.L. contributed to RNA-seq analysis. H.W. provided the anti-CD3 monoclonal antibody used in this study. H.W., I.A., B.L., M.M.., L.C. and J.L. contributed to the discussion and manuscript editing.

---

**Supplementary figure 1:**
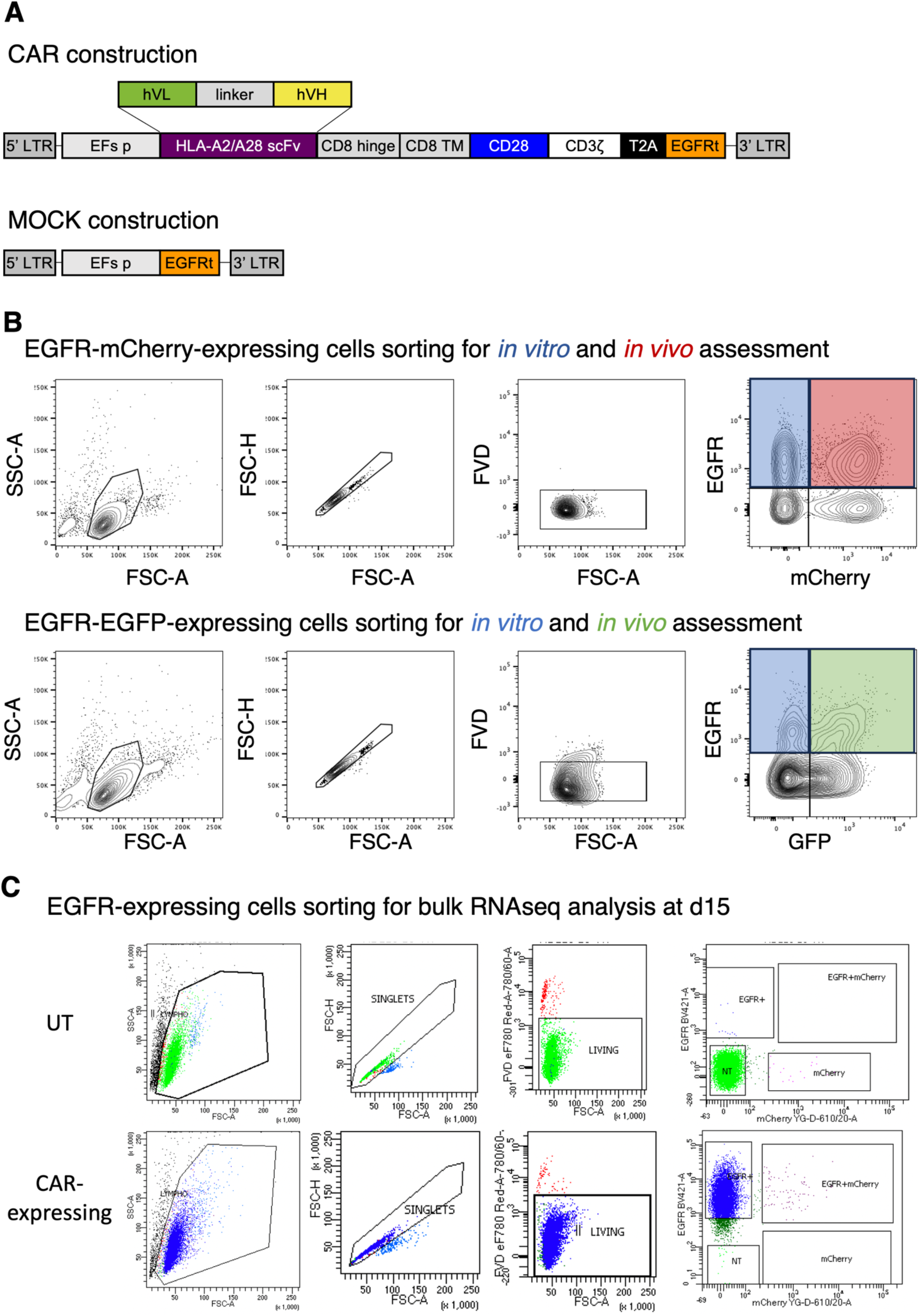
CAR construct and gating strategies used for CAR- and Luciferase- expressing cells sorting. **A.** CAR constructs against HLA-A2 and HLA-A28 and mock construct. **B.** (Up panel) Gating strategy to sort CAR-expressing (EGFR+) Tregs for *in vitro* assessment or CAR/Firefly Luciferase-expressing Tregs (EGFR+mCherry+) for *in vivo* assessment. (Down panel) Gating strategy to sort CAR-expressing (EGFR+) for *in vitro* assessment or CAR/CBG Luciferase-expressing (EGFR+GFP+) for *in vivo* assessment. **C.** Sorting strategy used for CAR-expressing cells isolation before bulk RNA sequencing.

**Supplementary figure 2:**
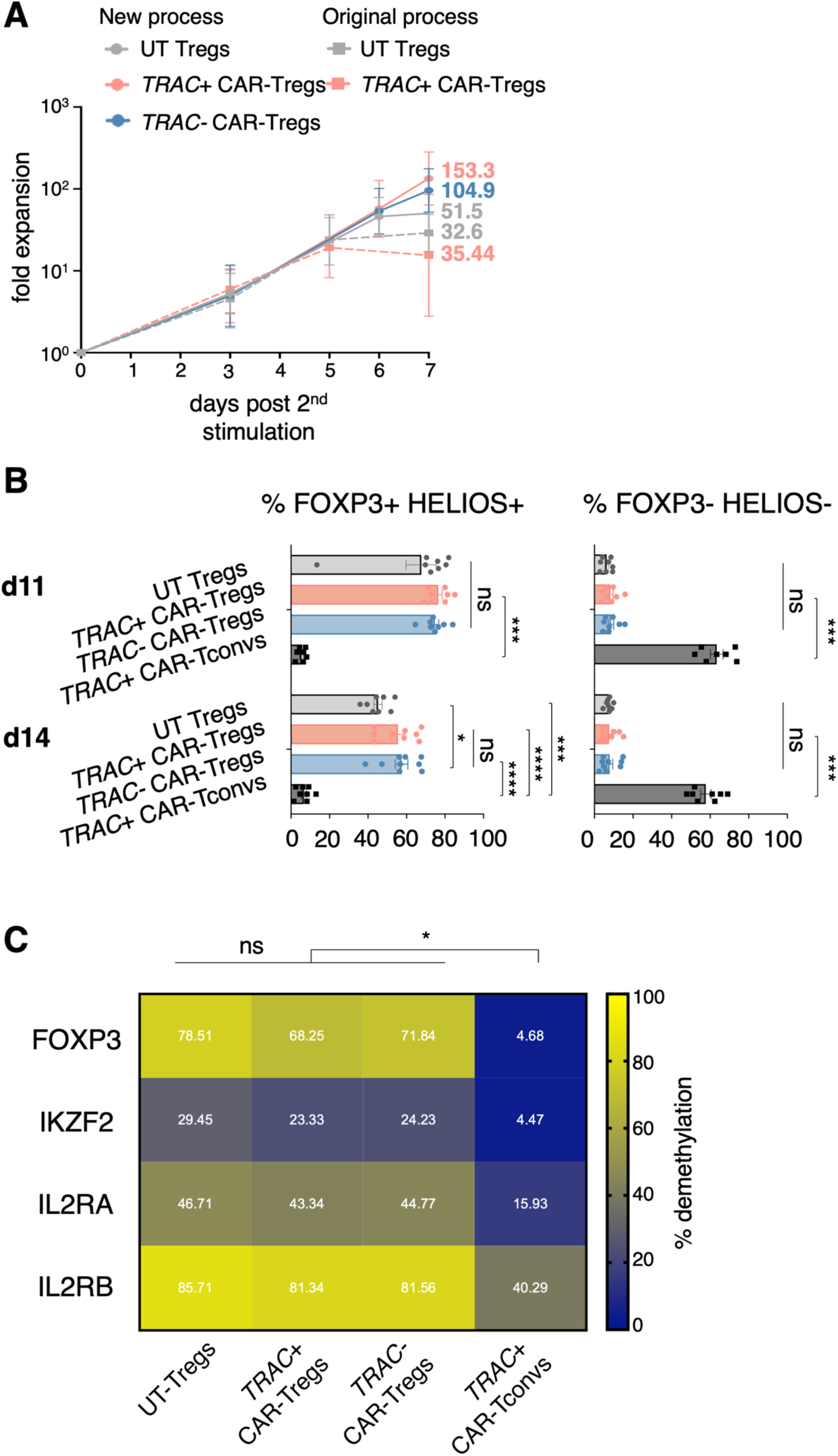
*TRAC-/-* CAR-Tregs maintain normal expansion, phenotypic and epigenetic stability upon CAR activation. **A.** Fold expansion of UT Tregs and CAR-Tregs (either *TRAC-/-* or *TRAC+/+*) following second stimulation for Tregs populations produced either with the new or original process^5^. The mean ± SD is shown. A total of 10 independent experiments are shown including the following cell populations: UT Tregs (grey full circle), *TRAC+/+* CAR-Tregs (red full circle) and *TRAC-/-* CAR-Tregs (blue full circle). Two-tailed Mann–Whitney indicated no significant differences in expansion between UT and CAR-Tregs at any timepoints. **B.** Frequencies of FOXP3+HELIOS+ double positive cells (left panel) and FOXP3-HELIOS- double negative cells (right panel) from at least 7 independent experiments. The mean ± SD is shown. All Tregs populations have a negligible proportion of double negative cells compared to CAR-Tconvs (p = 0.0003) on day 11 and (p < 0.0001) on day 15. **B.** bisulfite-conversion-based methylation PCR analysis on day 15 showing the mean percent demethylation of all CpG of the Treg-specific demethylated region (TSDR), IKZF2, IL2RA and IL2RB of UT Tregs, *TRAC-/-* CAR-Tregs, *TRAC+/+* CAR-Tregs and CAR-Tconvs from 4 independent experiments. A two-tailed Mann–Whitney test showed significant demethylation of the TSDR (p = 0.029), IKZF2 (p = 0.029), IL2RA (p = 0.029) and ILRRB (p = 0.029) compared to CAR-Tconvs.

**Supplementary figure 3:**
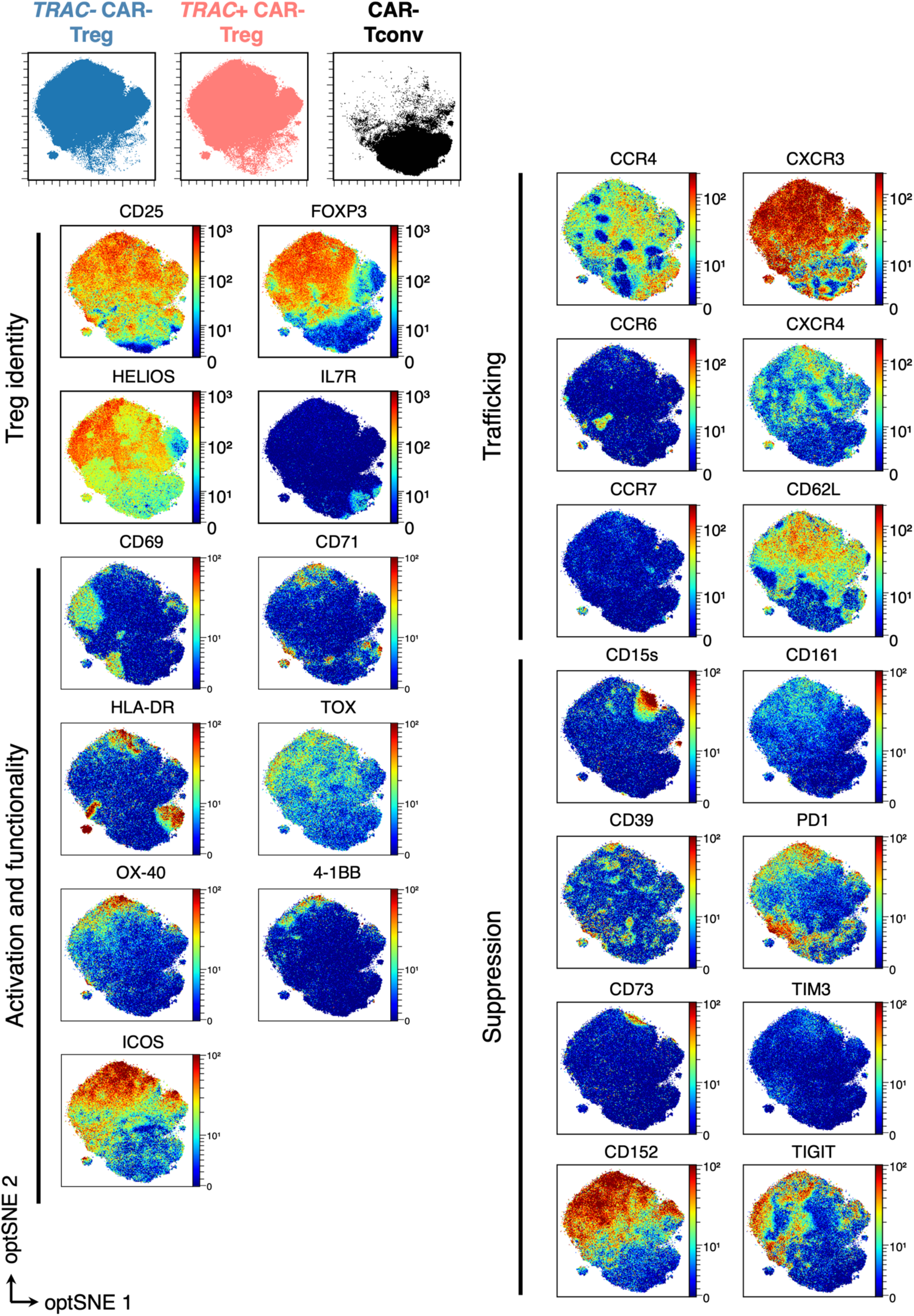
*TRAC*-/- CAR-Tregs and *TRAC*+/+ CAR-Tregs have similar Treg identity, activation and functionality, trafficking and suppression protein expression. Protein expression assessed by high-dimensional phenotypic analysis by Cytometry by Time-Of-Flight (CyTOF, mass cytometry) on day 15 of culture. Merged samples of 3 different donors. **A.** (Up panel) Spatial distribution of each cell population visualized by 2D opt-SNE: *TRAC*+/+ CAR-Tregs (red), *TRAC*-/- CAR-Tregs (blue), and CAR-Tconvs (black). (Bottom panel) Protein expression of key Treg markers projected onto the same 2D opt-SNE density map.

**Supplementary figure 4:**
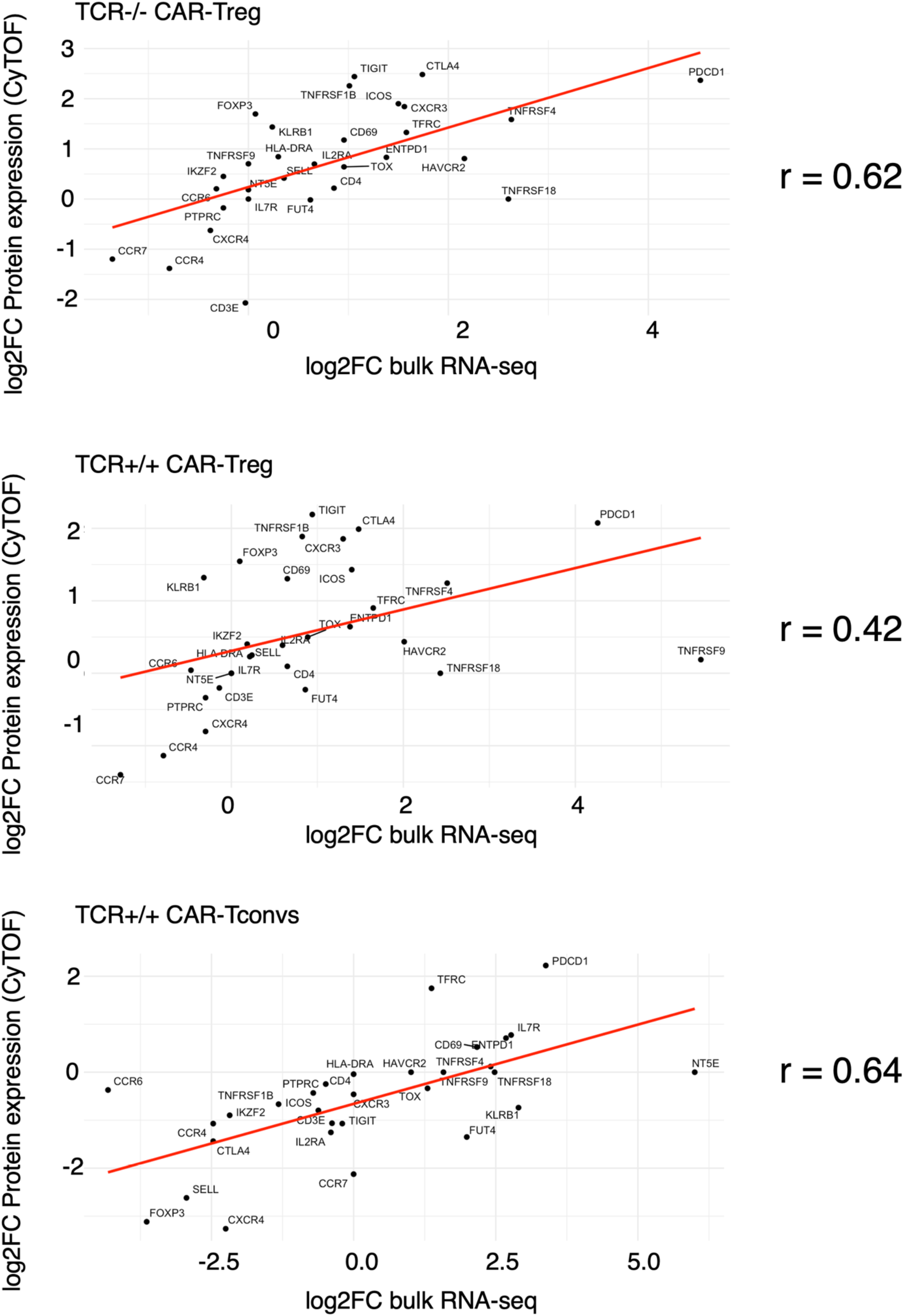
Transcriptomic and proteomic data have a good correlation across all conditions. Correlation curves comparing log₂ fold change relative to the UT Tregs condition between bulk RNA sequencing and CyTOF protein expression. Correlation coefficient for *TRAC*-/- CAR-Tregs, *TRAC*+/+ CAR-Tregs and CAR-Tconvs are displayed in the right panel.

**Supplementary figure 5:**
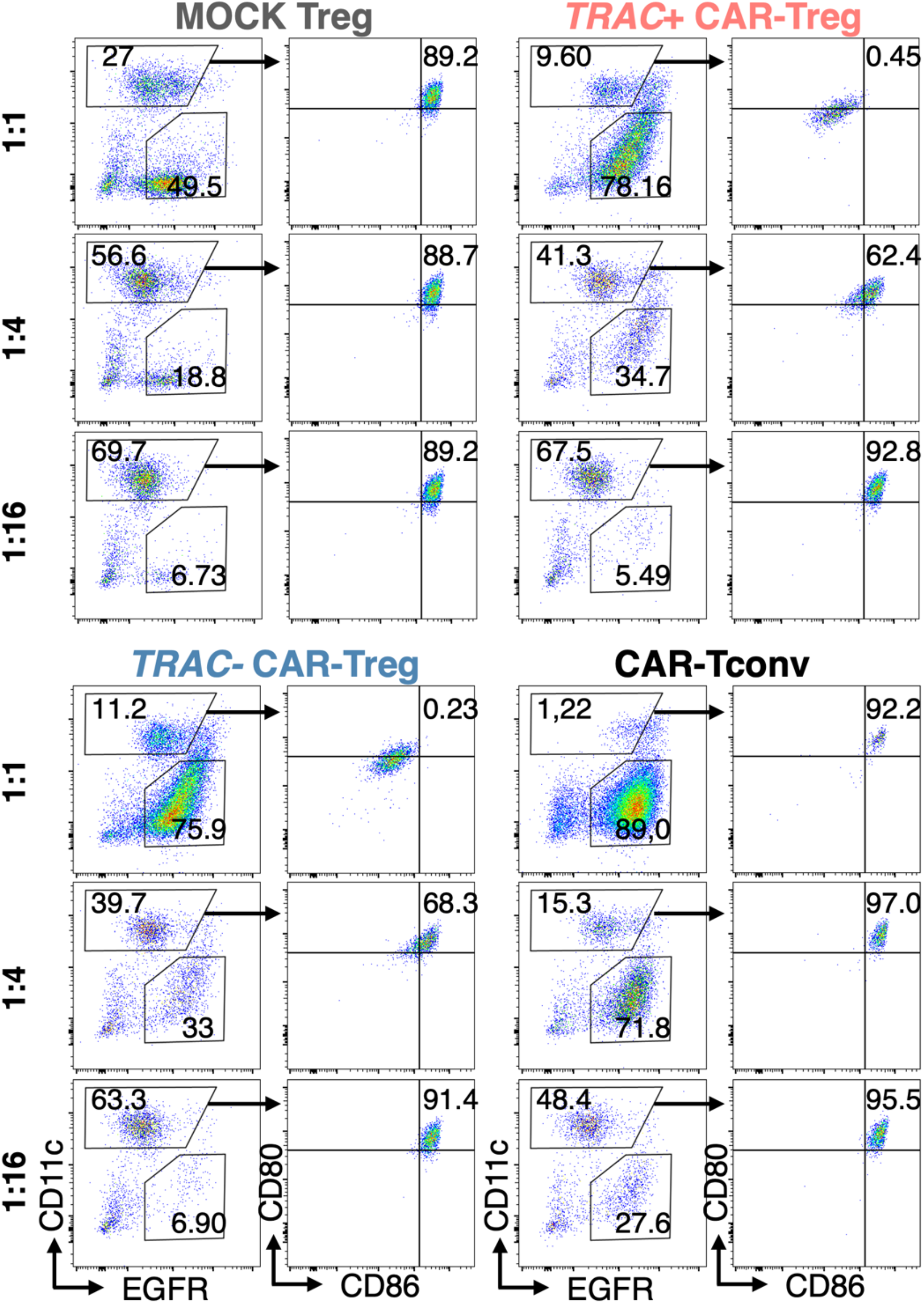
On day 16 of culture, MOCK Tregs, CAR-Tregs (either *TRAC-/-* or *TRAC+/+*) and CAR-Tconvs were put in co-culture with mature monocyte derived dendritic cells (moDC) during 40 hours with at the following T cell:moDC ratios: 1:1, 1:4, 1:16. **A.** Representative dot plots displaying CD11c (moDC) and EGFR (CAR- or MOCK-expressing cells) (left) and CD80/CD86 expression at the surface of moDC (right) for each co-culture: with MOCK Tregs (grey), *TRAC*+/+ CAR-Tregs (red), *TRAC*-/- CAR-Tregs (blue), and CAR-Tconvs (black).

**Supplementary figure 6:**
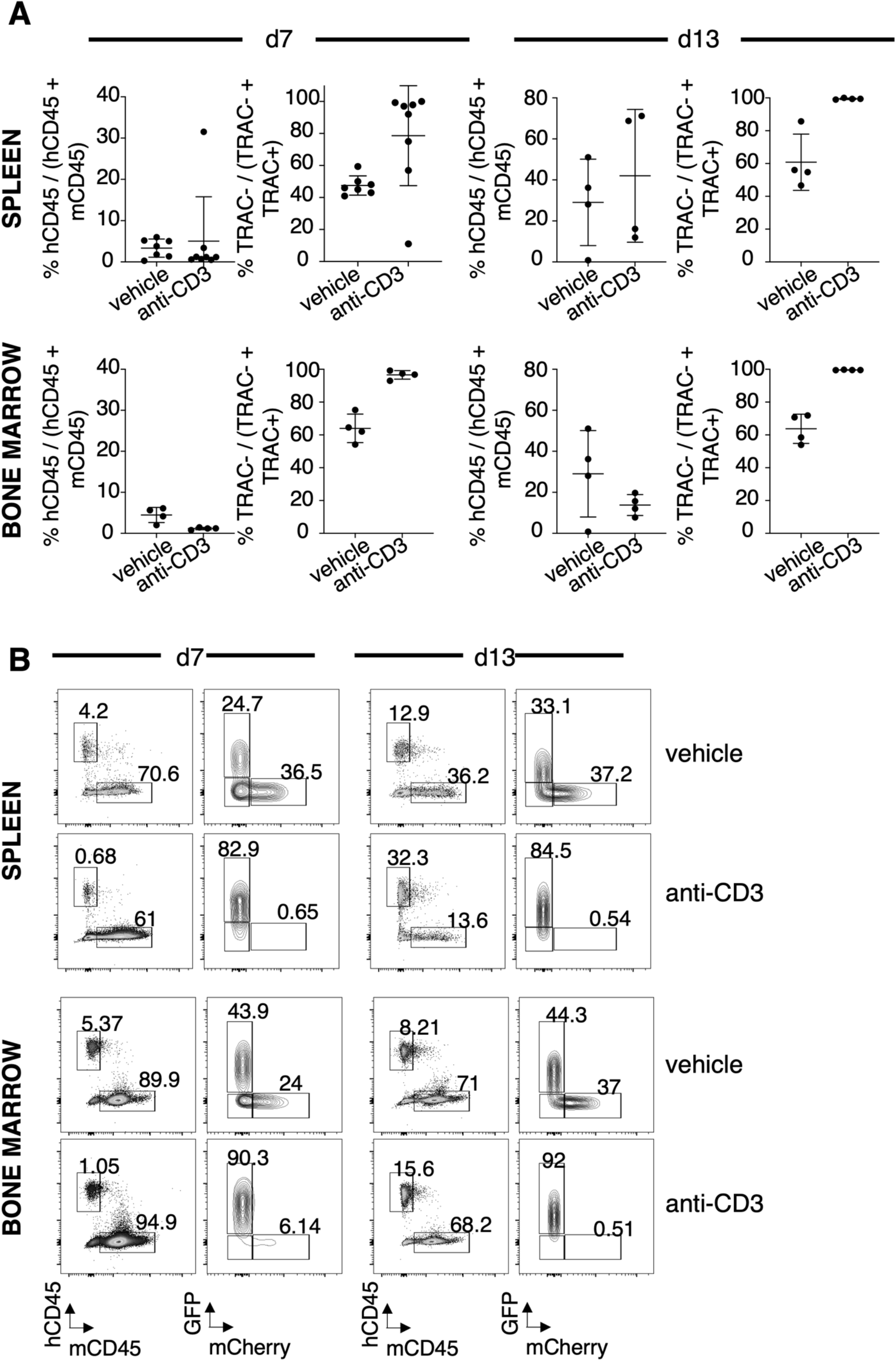
Non mitogenic anti-CD3 monoclonal antibody selectively targets *TRAC*+/+ conventional T cells in spleen and bone marrow. 8- to 12-week-old male NSG mice were conditioned with busulfan on days −2 and −1 before IV injection of 5x10^6^ HLA-A2+ PBMCs followed by retroorbital injection of 5×10^6^ total CAR-Tconvs (1:1 mixture of HLA-A2-targeted *TRAC*-/- GFP+ CAR-Tconvs and *TRAC*+/+ mCherry+ Luc+ CAR-Tconvs). Mice received 4 doses of 5 µg anti-CD3 on days 3, 4, 5 and 6 or were left untreated. **A-B.** Data from days 7 and 13, pooled from two independent experiments, are presented as mean ± SD. **A.** Quantification of human chimerism and the proportion of *TRAC*-/- cells among total CAR-Tconv populations in the spleen and bone marrow. **B.** Representative dot plots showing human CD45 versus murine CD45 expression and the distribution of GFP+ and mCherry+ cells within the human CD45+ population on spleen and bone marrow.

**Supplementary figure 7:**
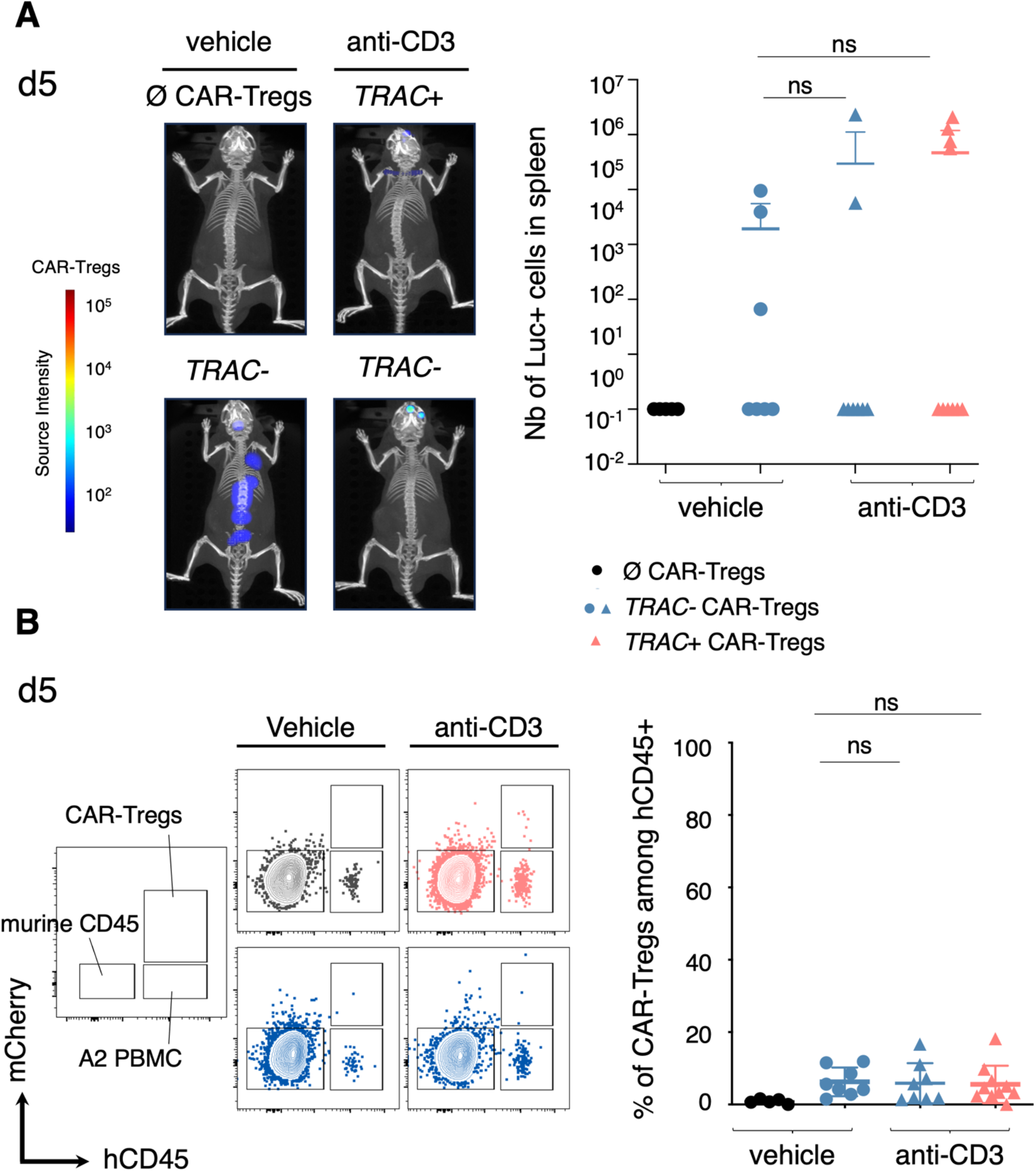
Non mitogenic anti-CD3 monoclonal antibody selectively targets *TRAC*+/+ regulatory T cells but spares their *TRAC*-/- counterpart. 8- to 12-week-old male NSG mice were conditioned with busulfan on days −2 and −1 before IV injection of 5x10^6^ HLA-A2+ PBMCs followed by retroorbital injection of 1×10^6^ Luc+ mCherry+ CAR-Tregs, either *TRAC*+/+ or *TRAC*-/-. Mice received 3 doses of 3 µg anti-CD3 on days 5, 6 and 7 or were left untreated (at least n=8 mice per group). **B-C.** Day 5 (before anti-CD3 treatment) data from 2 independent experiments are shown as mean ± SD. **B.** Representative 3D images (left panel) and total number of Luc+ cells was quantified by DLIT acquisition (right panel). A two-tailed Mann–Whitney test revealed no significant reduction in the number of *TRAC*+/+ CAR-Tregs (p=0.57) and *TRAC*-/- CAR-Tregs (p=0.65). **C.** Representative dot plots showing mCherry+ and human CD45+ cells (left panel) and quantification of the proportion of CAR-Tregs (*TRAC*+/+ or *TRAC*-/-) among human CD45 cells in peripheral blood (right panel). A two-tailed Mann–Whitney test showed no significant difference in the proportion of *TRAC*+/+ CAR-Tregs in anti-CD3-treated animals compared to *TRAC*-/- CAR- Tregs (p=0.55).

**Supplementary figure 8:**
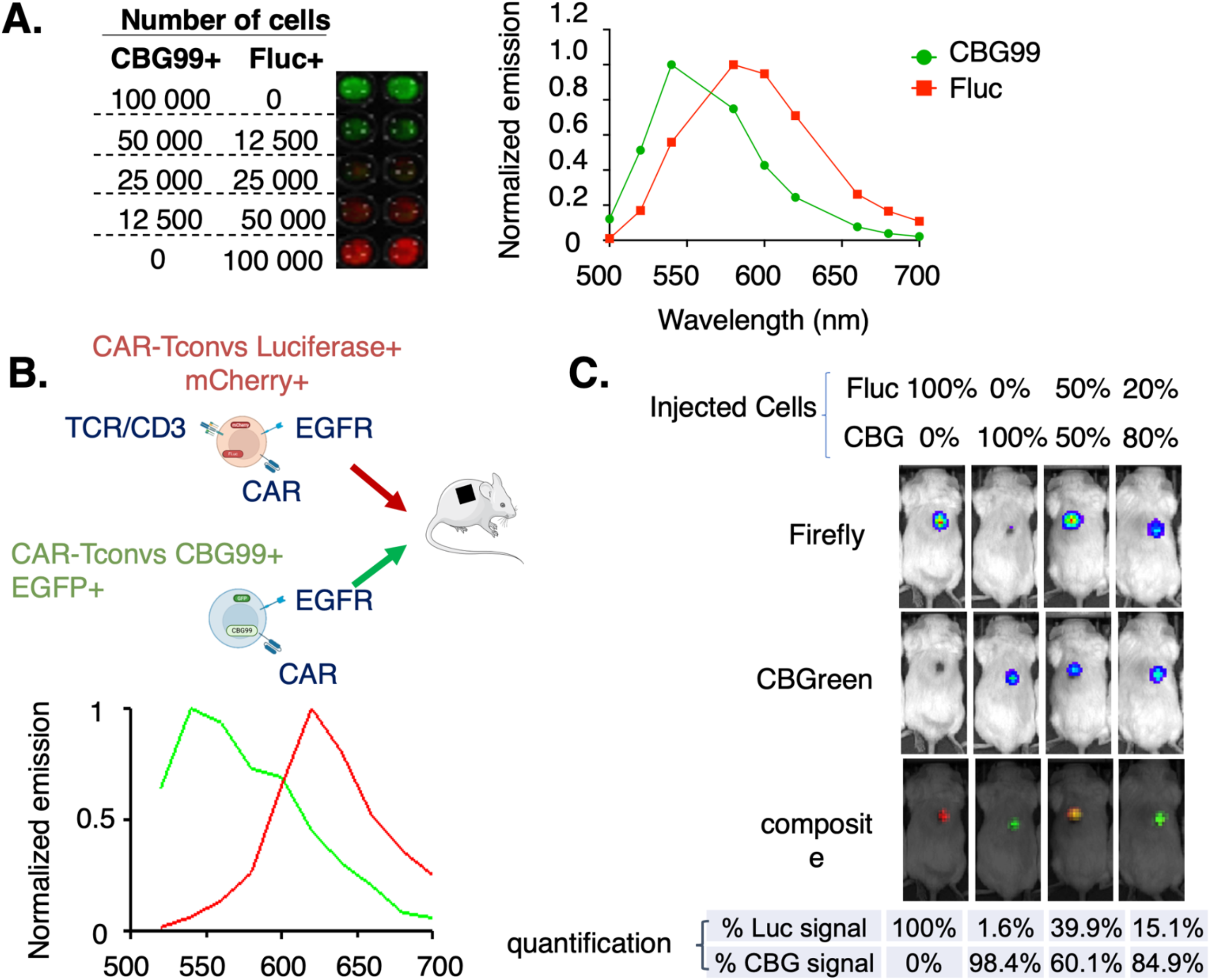
*In vitro* and *in vivo* dual luciferase acquisition optimization using spectral unmixing. **A.** (left panel) CD4+ T cells were transduced with either a Click Beetle Green (CBG) or Firefly (Fluc) luciferase, plated at varying ratios in a black 96-well plate, and imaged using the spectral unmixing plugin of the IVIS Spectrum™ system. (Right panel) *In vitro* spectral emission curves for CBG or Fluc luciferases are shown. **B-C.** NSG mice were grafted with HLA-A2-expressing allogeneic skins prior to administration of CBG- or Fluc-expressing CAR-T cells. **B.** Schematic representation of the experimental design and *in vivo* spectral emission of CBG or Fluc luciferases. **C.** Skin-grafted mice were infused with varying ratios of CBG- or Fluc-expressing CAR-Tconvs. The injected cell proportions correspond to the ratios quantified on day 19 post-injection using spectral unmixing.

**Supplementary figure 9:**
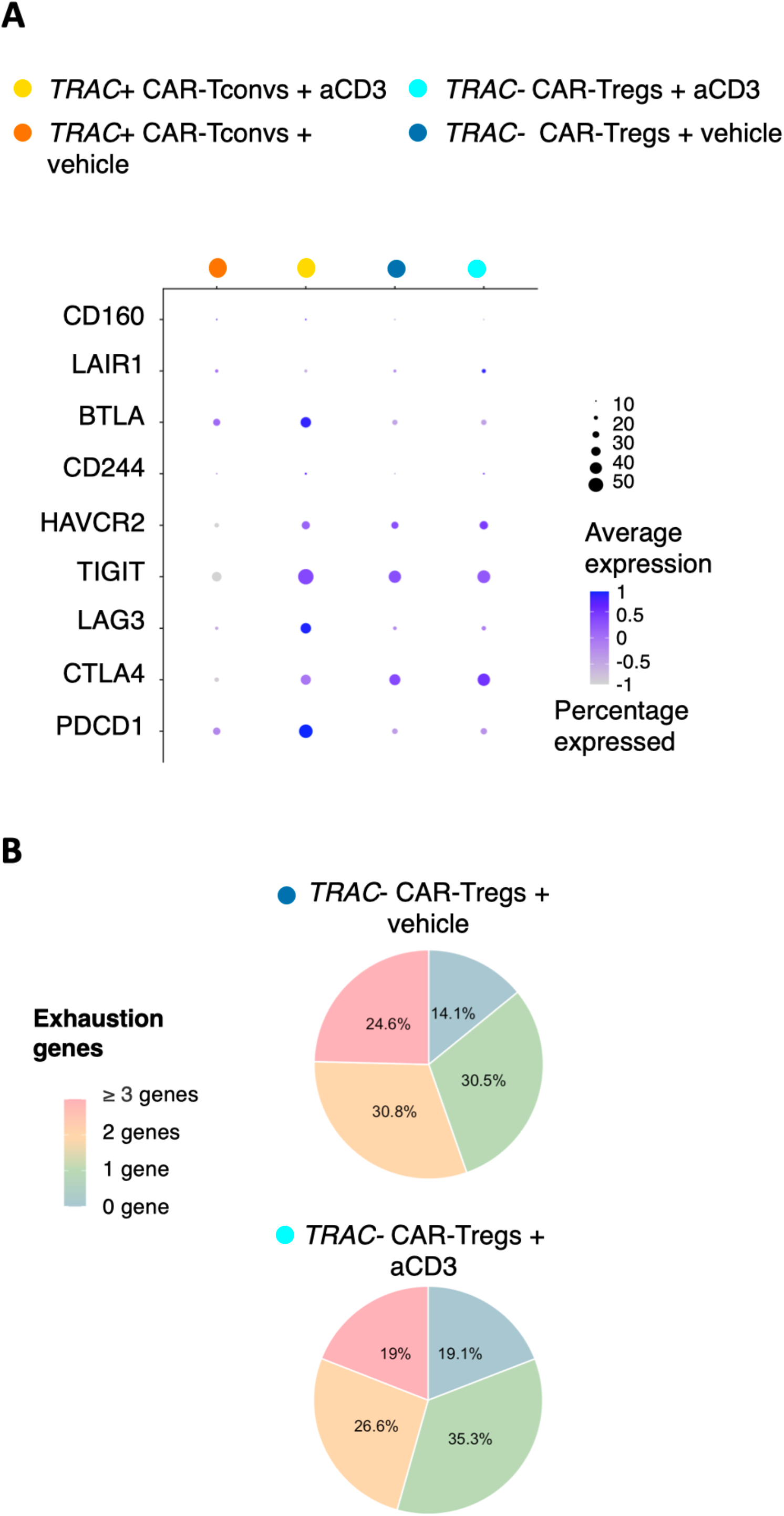
*TRAC*-/- CAR-Tregs maintain their transcriptomic profile following anti-CD3 treatment. *TRAC*+/+ CAR-Tconvs and *TRAC*-/- CAR-Tregs were collected 12 hours after anti-CD3 administration from the spleens of 8- to 12- week-old preconditioned NSG male mice injected with HLA-A2+ PBMC and a 1:2 ratio of *TRAC*+/+ CAR-Tconvs to *TRAC*-/- CAR-Tregs. **A.** Single-cell RNA-seq analysis from one experiment, including 3 mice per group **A.** Dot plots generated using the Seurat package displaying the expression of selected exhaustion-related genes. **B.** Pie charts summarizing the proportion of cells expressing 0, 1 or co-expressing 2 or ≥3 exhaustion genes, comparing TRAC-/- CAR-Tregs from anti-CD3-treated vs untreated mice.

**Supplementary figure 10:**
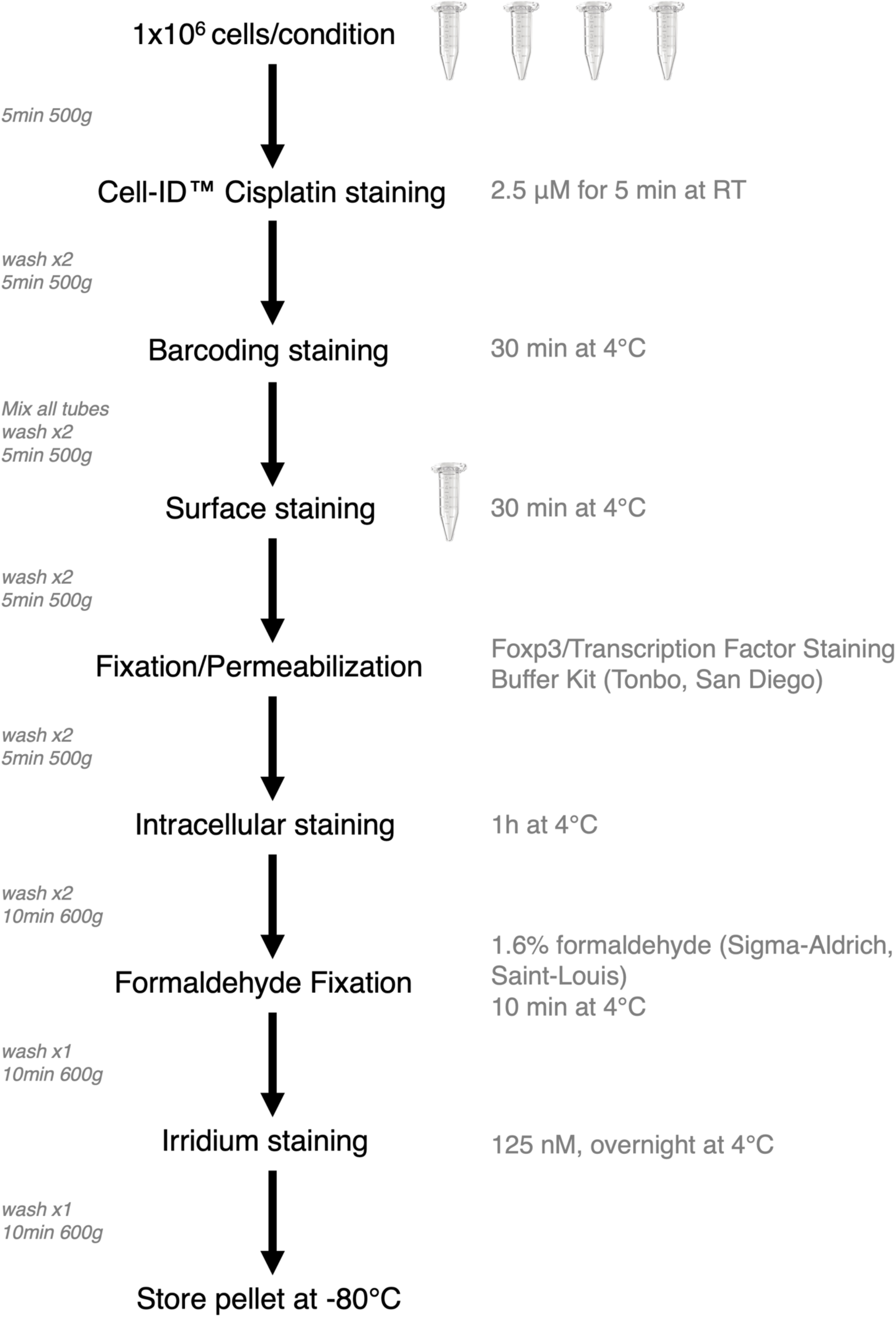
Schematic representation of the mass spectrometry staining procedure for high dimensional cell characterization.

**SUPPLEMENTARY TABLE 1:**
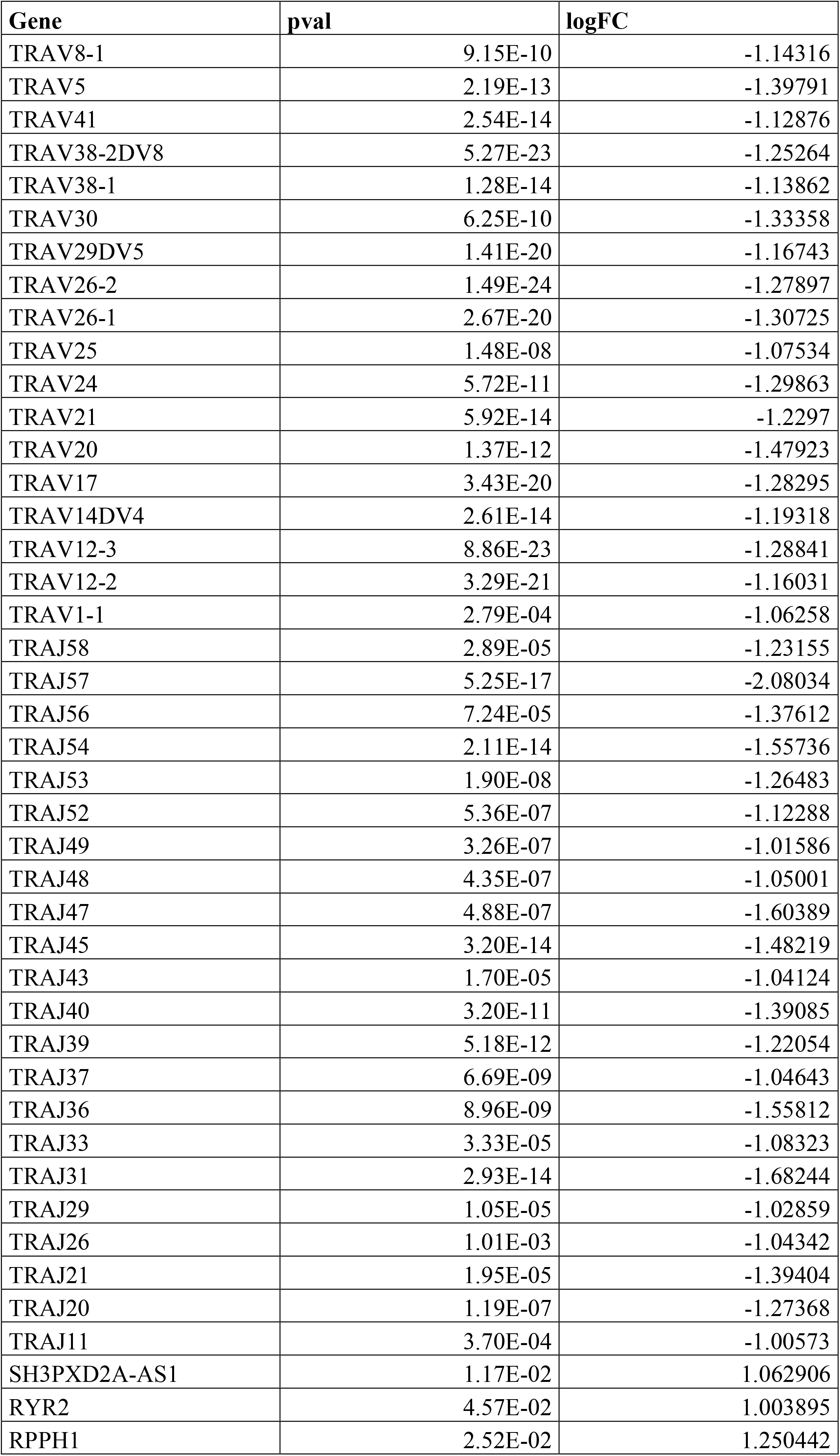
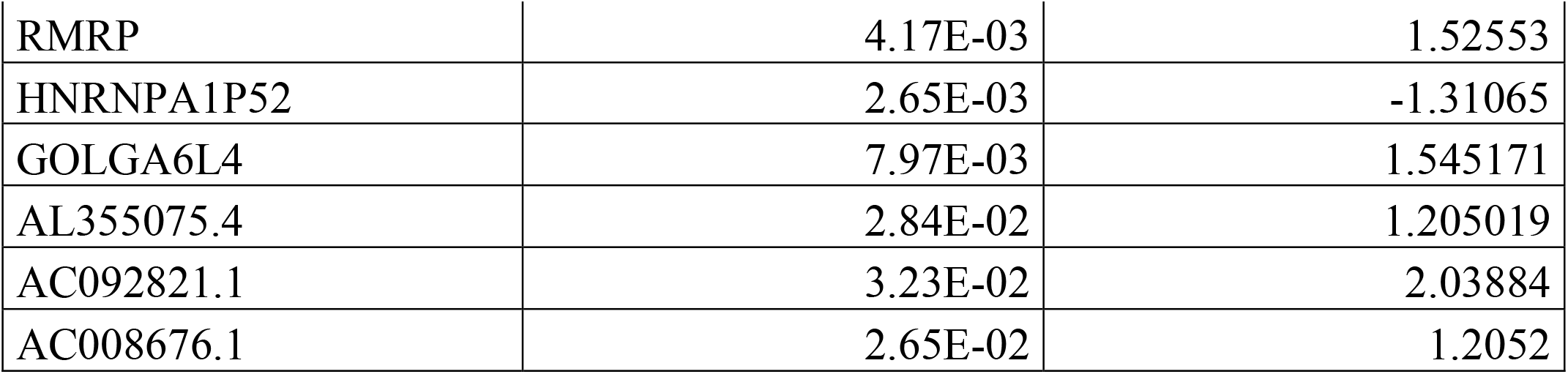
List of differentially expressed genes comparing *TRAC*-/- CAR-Tregs vs *TRAC*+/+ CAR-Tregs.

**SUPPLEMENTARY TABLE 2:** List of antibodies used for high-dimensional phenotypic analysis by Cytometry by Time-Of-Flight (CyTOF, mass cytometry)

| Name | Fluorophore | Clone | Manufacturer | Catalog number | Dilution | Panel |
| --- | --- | --- | --- | --- | --- | --- |
| human CD45 | 116Cd | HI30 | Standard Bitools | 3116001C | 1µL/5M | Cytof barcoding |
| human CD45 | 110CD | HI30 | Standard Bitools | 3110001C | 1µL/5M |  |
| human CD45 | 112CD | HI30 | Standard Bitools | 3112001C | 1µL/5M |  |
| human CD45 | 106Cd | HI30 | Standard Bitools | 3106001C | 1µL/5M |  |
| CD69 | 113Cd | FN50 | Standard Bitools | 3113002C | 1µL/5M | Cytof surface markers |
| CD3 | 141Pr | UCHT1 | Standard Bitools | 3141019C | 0.5µL/5M |  |
| CD45RA | 143Nd | HI100 | Standard Bitools | 3143006C | 1µL/5M |  |
| CD4 | 145Nd | SK3 | Standard Bitools | 92J033145 | 0.5µL/5M |  |
| TNFRII | 147Sm | Polyclonal | Biotechne | AF726 | 2µL/5M |  |
| CD278 / ICOS | 148Nd | C398.4A | Standard Bitools | 3148019C | 2µL/5M |  |
| CCR4 | 149Sm | L291H4 | Standard Bitools | 3149029C | 2µL/5M |  |
| CD15s | 151Eu | CLEX1 | BD Biosciences | 551344 | 2µL/5M |  |
| TIM-3 | 153Eu | F38-2E2 | Standard Bitools | 3153008C | 2µL/5M |  |
| TIGIT | 154Sm | MBSA43 | Standard Bitools | 3154016C | 2µL/5M |  |
| PD-1 | 155Gd | EH12.2H7 | Standard Bitools | 3155009C | 2µL/5M |  |
| CD134 | 158Gd | ACT35 | Standard Bitools | 3158012C | 2µL/5M |  |
| CD357/GITR | 159Tb | 621 | Standard Bitools | 3159020C | 2µL/5M |  |
| CD39 | 160Gd | A1 | Standard Bitools | 3160004C | 2µL/5M |  |
| CD183 | 163Dy | G025H7 | Standard Bitools | 3163004C | 2µL/5M |  |
| CD161 | 164Dy | HP-3G10 | Standard Bitools | 3164009C | 2µL/5M |  |
| CD127 | 165Ho | A019D5 | Standard Bitools | 3165008C | 1µL/5M |  |
| CD197 | 167Er | G043H7 | Standard Bitools | 3167009C | 2µL/5M |  |
| CD73 | 168Er | AD2 | Standard Bitools | 3168015C | 2µL/5M |  |
| CD25 | 169Tm | 2A3 | Standard Bitools | 3169003C | 1µL/5M |  |
| CXCR4 | 173Yb | 12G5 | Standard Bitools | 3173001C | 2µL/5M |  |
| HLA-DR | 174Yb | L243 | Standard Bitools | 3174001C | 2µL/5M |  |
| CD71 | 175Lu | OKT-9 | Standard Bitools | 3175011C | 2µL/5M |  |
| CD196/CCR6 | 176Yb | G034E3 | Standard Bitools | 3176022C | 1µL/5M |  |
| CD62L | 171Yb | DREG-56 | BD Biosciences | 555542 | 2µL/5M |  |
| CD137 | 209Bi | 4B4-1 | Standard Bitools | 3209015C | 2µL/5M |  |
| EGFR | 170Er | AY13 | Standard Bitools | 3170009B | 2µL/5M |  |
| CD152 | 142Nd | 14D3 | Standard Bitools | 3142019C | 2µL/5M | Cytof intracellular markers |
| Helios | 156Gd | 22F6 | Biologend | 137202 | 1µL/5M |  |
| FOXP3 | 162Dy | PCH101 | Standard Bitools | 3162011C | 2µL/5M |  |
| TOX | 172Yb | TXRX10 | ThermoFisher Scientific | 14-6502-82 | 2µL/5M |  |

**SUPPLEMENTARY TABLE 3:**
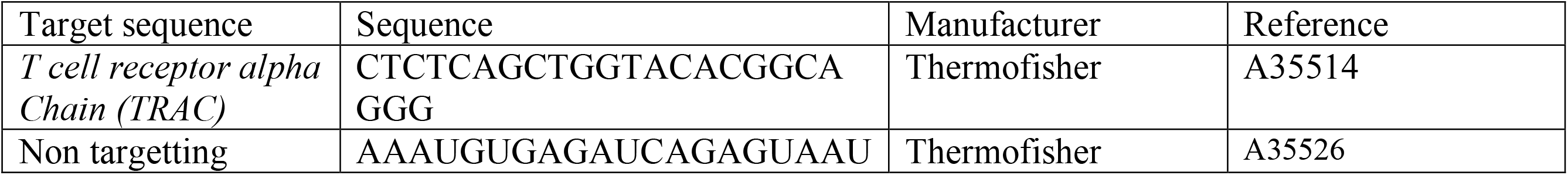
sgRNA used for gene editing.

**SUPPLEMENTARY TABLE 4:** List of antibodies and staining reagents.

| Name | Fluorophore | Clone | Manufacturer | Catalog number | Dilution | Panel |
| --- | --- | --- | --- | --- | --- | --- |
| 7-AAD | - | - | Miltenyi Biotec | 130-111-568 | 1/100 | Naive Treg sorting |
| CD127 | Brillant Violet 421 | HIL-7R-M21 | BD Biosciences | 562436 | 1/50 |  |
| CD25 | PE | M-A251 | BD Biosciences | 555432 | 1/10 |  |
| CD4 | FITC | SK3/SK4 | BD Biosciences | 347413 | 1/5 |  |
| CD45RA | Brillant Violet 711 | HI100 | Ozyme | 304138 | 1/50 |  |
| CD45RO | APC | UCHL1 | BD Biosciences | 559865 | 1/25 |  |
| EGFR | Brillant Violet 421 | AY13 | Biolegend | 352911 | 1/50 | Phenotyping |
| TCR alpha/beta | FITC | BW242/412 | Miltenyi Biotec | 130-098-688 | 1/50 |  |
| CD3 | Brillant Violet 510 | UCHT-1 | BD Biosciences | 563109 | 1/20 |  |
| HELIOS | PE-Cy7 | 22F6 | ThermoFisher Scientific | 25-9883-42 | 1/20 |  |
| FOXP3 | PerCP Cy5.5 | 236A/E7 | BD Bioscience | 561493 | 1/20 |  |
| FVD eF780 | eFluor 780 |  | ThermoFisher Scientific | 65-0865-18 | 1/1000 |  |
| CD4 | PerCP Cy5.5 | RPA-T4 | BD Biosciences | 560650 | 1/20 | Activation assay |
| GARP | Brillant Violet 711 | 7B11 | BD Biosciences | 563958 | 1/20 |  |
| CD137 | Brillant Violet 605 | 4B4-1 | Sony biotechnology | 2149140 | 1/20 |  |
| CD69 | eFluor 450 | FN50 | ThermoFisher Scientific | 48-0699-42 | 1/28 |  |
| HLA-DR | PE | G46-6 | BD Biosciences | 555812 | 1/5 |  |
| CD25 | Brillant Violet 510 | M-A251 | BD Biosciences | 563352 | 1/20 |  |
| CD11c | PE-Cy7 | B-ly6 | BD Biosciences | 561356 | 1/20 | Transendocytosis assay |
| CD80 | BV510 | 2D10 | Biolegend | 305233 | 1/20 |  |
| CD86 | PE | BU63 | Biolegend | 374205 | 1/20 |  |
| mouse CD45 | PE-Cy7 | 30-F11 | Sony biotechnology | 1115570 | 1/20 | *in vivo* chimerism |
| human CD45 | Brillant Violet 421 | HI30 | BD Biosciences | 563879 | 1/20 |  |

## REFERENCES

1. Sagoo, P., Ali, N., Garg, G., Nestle, F.O., Lechler, R.I., and Lombardi, G. (2011). Human regulatory T cells with alloantigen specificity are more potent inhibitors of alloimmune skin graft damage than polyclonal regulatory T cells. Sci. Transl. Med. 3. 10.1126/SCITRANSLMED.3002076,.

2. Sawitzki, B., Harden, P.N., Reinke, P., Moreau, A., Hutchinson, J.A., Game, D.S., Tang, Q., Guinan, E.C., Battaglia, M., Burlingham, W.J., et al. (2020). Regulatory cell therapy in kidney transplantation (The ONE Study): a harmonised design and analysis of seven non-randomised, single-arm, phase 1/2A trials. The Lancet 395, 1627–1639. 10.1016/S0140-6736(20)30167-7.

3. Tang, Q., Leung, J., Peng, Y., Sanchez-Fueyo, A., Lozano, J.J., Lam, A., Lee, K., Greenland, J.R., Hellerstein, M., Fitch, M., et al. (2022). Selective decrease of donor-reactive Tregs after liver transplantation limits Treg therapy for promoting allograft tolerance in humans. Sci. Transl. Med. 14. 10.1126/SCITRANSLMED.ABO2628,.

4. Dawson, N.A.J., Rosado-Sánchez, I., Novakovsky, G.E., Fung, V.C.W., Huang, Q., McIver, E., Sun, G., Gillies, J., Speck, M., Orban, P.C., et al. (2020). Functional effects of chimeric antigen receptor co-receptor signaling domains in human regulatory T cells. Sci. Transl. Med. 12. 10.1126/SCITRANSLMED.AAZ3866,.

5. Lamarthée, B., Marchal, A., Charbonnier, S., Blein, T., Leon, J., Martin, E., Rabaux, L., Vogt, K., Titeux, M., Delville, M., et al. (2021). Transient mTOR inhibition rescues 4-1BB CAR-Tregs from tonic signal-induced dysfunction. Nat. Commun. 12. 10.1038/S41467-021-26844-1.

6. Kurt, A.S., Ruiz, P., Landmann, E., Elgosbi, M., Kan Fung, T., Kodela, E., Londoño, M.C., Correa, D.M., Perpiñán, E., Lombardi, G., et al. (2024). Conferring alloantigen specificity to regulatory T cells: A comparative analysis of cell preparations undergoing clinical development in transplantation. American Journal of Transplantation 25. 10.1016/j.ajt.2024.09.009.

7. Stark, H., Ho, Q.Y., Cross, A., Alessandrini, A., Bertaina, A., Brennan, D., Busque, S., Demetris, A., Devey, L., Fruhwirth, G., et al. (2025). Meeting Report: The Sixth International Sam Strober Workshop on Clinical Immune Tolerance. Transplantation 109. 10.1097/TP.0000000000005311,.

8. DeWolf, S., Grinshpun, B., Savage, T., Lau, S.P., Obradovic, A., Shonts, B., Yang, S., Morris, H., Zuber, J., Winchester, R., et al. (2018). Quantifying size and diversity of the human T cell alloresponse. JCI Insight 3. 10.1172/JCI.INSIGHT.121256,.

9. Beura, L.K., Hamilton, S.E., Bi, K., Schenkel, J.M., Odumade, O.A., Casey, K.A., Thompson, E.A., Fraser, K.A., Rosato, P.C., Filali-Mouhim, A., et al. (2016). Normalizing the environment recapitulates adult human immune traits in laboratory mice. Nature 532, 512–516. 10.1038/NATURE17655,.

10. Stranavova, L., Pelak, O., Svaton, M., Hruba, P., Fronkova, E., Slavcev, A., Osickova, K., Maluskova, J., Hubacek, P., Fronek, J., et al. (2019). Heterologous cytomegalovirus and allo-reactivity by shared t cell receptor repertoire in kidney transplantation. Front. Immunol. 10. 10.3389/FIMMU.2019.02549,.

11. Adams, A.B., Williams, M.A., Jones, T.R., Shirasugi, N., Durham, M.M., Kaech, S.M., Wherry, E.J., Onami, T., Lanier, J.G., Kokko, K.E., et al. (2003). Heterologous immunity provides a potent barrier to transplantation tolerance. Journal of Clinical Investigation 111, 1887–1895. 10.1172/JCI17477,.

12. Benichou, G., Gonzalez, B., Marino, J., Ayasoufi, K., and Valujskikh, A. (2017). Role of memory T cells in allograft rejection and tolerance. Front. Immunol. 8. 10.3389/FIMMU.2017.00170,.

13. Cravedi, P., Riella, L. V., Ford, M.L., Valujskikh, A., Menon, M.C., Kirk, A.D., Alegre, M.L., Alessandrini, A., Feng, S., Kehn, P., et al. (2024). Advancing mouse models for transplantation research. American Journal of Transplantation 24, 1362–1368. 10.1016/j.ajt.2024.01.006.

14. Todo, S., Yamashita, K., Goto, R., Zaitsu, M., Nagatsu, A., Oura, T., Watanabe, M., Aoyagi, T., Suzuki, T., Shimamura, T., et al. (2016). A pilot study of operational tolerance with a regulatory T-cell-based cell therapy in living donor liver transplantation. Hepatology 64, 632–643. 10.1002/HEP.28459,.

15. You, S., Zuber, J., Kuhn, C., Baas, M., Valette, F., Sauvaget, V., Sarnacki, S., Sawitzki, B., Bach, J.F., Volk, H.D., et al. (2012). Induction of allograft tolerance by monoclonal CD3 antibodies: A matter of timing. American Journal of Transplantation 12, 2909–2919. 10.1111/j.1600-6143.2012.04213.x.

16. Goto, R., You, S., Zaitsu, M., Chatenoud, L., and Wood, K.J. (2013). Delayed anti-CD3 therapy results in depletion of alloreactive T cells and the dominance of foxp3+Cd4+ graft infiltrating cells. American Journal of Transplantation 13, 1655–1664. 10.1111/ajt.12272.

17. Perruche, S., Zhang, P., Liu, Y., Saas, P., Bluestone, J.A., and Chen, W.J. (2008). CD3-specific antibody-induced immune tolerance involves transforming growth factor-β from phagocytes digesting apoptotic T cells. Nat. Med. 14, 528–535. 10.1038/NM1749,.

18. Belghith, M., Bluestone, J.A., Barriot, S., Mégret, J., Bach, J.F., and Chatenoud, L. (2003). TGF-β-dependent mechanisms mediate restoration of self-tolerance induced by antibodies to CD3 in overt autoimmune diabetes. Nat. Med. 9, 1202– 1208. 10.1038/NM924,.

19. Dong, D., Zheng, L., Lin, J., Zhang, B., Zhu, Y., Li, N., Xie, S., Wang, Y., Gao, N., and Huang, Z. (2019). Structural basis of assembly of the human T cell receptor–CD3 complex. Nature 2019 573:7775 573, 546–552. 10.1038/S41586-019-1537-0.

20. Hoffmann, P., Boeld, T.J., Eder, R., Huehn, J., Floess, S., Wieczorek, G., Olek, S., Dietmaier, W., Andreesen, R., and Edinger, M. (2009). Loss of FOXP3 expression in natural human CD4+ CD25+ regulatory T cells upon repetitive in vitro stimulation. Eur. J. Immunol. 39, 1088–1097. 10.1002/EJI.200838904,.

21. Feng, Y., Arvey, A., Chinen, T., Van Der Veeken, J., Gasteiger, G., and Rudensky, A.Y. (2014). Control of the inheritance of regulatory T cell identity by a cis element in the foxp3 locus. Cell 158, 749–763. 10.1016/j.cell.2014.07.031.

22. Pesenacker, A.M., Wang, A.Y., Singh, A., Gillies, J., Kim, Y., Piccirillo, C.A., Nguyen, D., Haining, W.N., Tebbutt, S.J., Panagiotopoulos, C., et al. (2016). A Regulatory T-Cell Gene Signature Is a Specific and Sensitive Biomarker to Identify Children With New-Onset Type 1 Diabetes. Diabetes 65, 1031–1039. 10.2337/DB15-0572.

23. Tekguc, M., Wing, J.B., Osaki, M., Long, J., and Sakaguchi, S. (2021). Treg-expressed CTLA-4 depletes CD80/CD86 by trogocytosis, releasing free PD-L1 on antigen-presenting cells. Proc. Natl. Acad. Sci. U. S. A. 118. 10.1073/PNAS.2023739118.

24. Bolt, S., Routledge, E., Lloyd, I., Chatenoud, L., Pope, H., Gorman, S.D., Clark, M., and Waldmann, H. (1993). The generation of a humanized, non-mitogenic CD3 monoclonal antibody which retains in vitro immunosuppressive properties. Eur. J. Immunol. 23, 403–411. 10.1002/EJI.1830230216.

25. Friend, P.J., Hale, G., Chatenoud, L., Rebello, P., Bradley, J., Thiru, S., Phillips, J.M., and Waldmann, H. (1999). Phase I study of an engineered aglycosylated humanized CD3 antibody in renal transplant rejection. Transplantation 68, 1632– 1637. 10.1097/00007890-199912150-00005.

26. Li, Y., Li, X.C., Zheng, X.X., Wells, A.D., Turka, L.A., and Strom, T.B. (1999). Blocking both signal 1 and signal 2 of T-cell activation prevents apoptosis of alloreactive t cells and induction of peripheral allograft tolerance. Nat. Med. 5, 1298–1302. 10.1038/15256,.

27. Zheng, X.X., Sánchez-Fueyo, A., Sho, M., Domenig, C., Sayegh, M.H., and Strom, T.B. (2003). Favorably tipping the balance between cytopathic and regulatory T cells to create transplantation tolerance. Immunity 19, 503–514. 10.1016/S1074-7613(03)00259-0.

28. Picarda, E., Bézie, S., Boucault, L., Autrusseau, E., Kilens, S., Meistermann, D., Martinet, B., Daguin, V., Donnart, A., Charpentier, E., et al. (2017). Transient antibody targeting of CD45RC induces transplant tolerance and potent antigen-specific regulatory T cells. JCI Insight 2. 10.1172/JCI.INSIGHT.90088,.

29. Blein, T., Ayas, N., Charbonnier, S., Léon, J., and Zuber, J. (2025). Tolerance Induction Strategies in Organ Transplantation: Current Status and Future Perspectives. manuscript under revision by Transplant International.

30. Yamada, Y., Boskovic, S., Aoyama, A., Murakami, T., Putheti, P., Smith, R.N., Ochiai, T., Nadazdin, O., Koyama, I., Boenisch, O., et al. (2012). Overcoming memory T-cell responses for induction of delayed tolerance in nonhuman primates. American Journal of Transplantation 12, 330–340. 10.1111/j.1600-6143.2011.03795.x.

31. Keymeulen, B., Vandemeulebroucke, E., Ziegler, A.G., Mathieu, C., Kaufman, L., Hale, G., Gorus, F., Goldman, M., Walter, M., Candon, S., et al. (2005). Insulin Needs after CD3-Antibody Therapy in New-Onset Type 1 Diabetes. New England Journal of Medicine 352, 2598–2608. 10.1056/NEJMOA043980/SUPPL_FILE/2598SA1-4.PDF.

32. Keymeulen, B., Walter, M., Mathieu, C., Kaufman, L., Gorus, F., Hilbrands, R., Vandemeulebroucke, E., Van De Velde, U., Crenier, L., De Block, C., et al. (2010). Four-year metabolic outcome of a randomised controlled CD3-antibody trial in recent-onset type 1 diabetic patients depends on their age and baseline residual beta cell mass. Diabetologia 53, 614–623. 10.1007/S00125-009-1644-9,.

33. Ramos, E.L., Dayan, C.M., Chatenoud, L., Sumnik, Z., Simmons, K.M., Szypowska, A., Gitelman, S.E., Knecht, L.A., Niemoeller, E., Tian, W., et al. (2023). Teplizumab and β-Cell Function in Newly Diagnosed Type 1 Diabetes. New England Journal of Medicine 389, 2151–2161. 10.1056/NEJMOA2308743,.

34. Vudattu, N.K., and Herold, K.C. (2013). Delayed anti-CD3 therapy in a mouse heart transplant model induced tolerance and long-term survival of allograft: Achieving tolerance. Immunotherapy 5, 1173–1176. 10.2217/IMT.13.113,.

35. Reinke, P., Schwinzer, H., Höflich, C., Ode-Hakim, C., Döcke, W.D., Frei, U., and Volk, H.D. (1997). Selective in vivo deletion of alloactivated TH1 cells by OKT3 monoclonal antibody in acute rejection. Immunol. Lett. 57, 151–153. 10.1016/S0165-2478(97)00092-8.

36. Waldron-Lynch, F., Henegariu, O., Deng, S., Preston-Hurlburt, P., Tooley, J., Flavell, R., and Herold, K.C. (2012). Teplizumab induces human gut-tropic regulatory cells in humanized mice and patients. Sci. Transl. Med. 4. 10.1126/SCITRANSLMED.3003401,.

37. Long, S.A., Thorpe, J., DeBerg, H.A., Gersuk, V., Eddy, J.A., Harris, K.M., Ehlers, M., Herold, K.C., Nepom, G.T., and Linsley, P.S. (2016). Partial exhaustion of CD8 T cells and clinical response to teplizumab in new-onset type 1 diabetes. Sci. Immunol. 1. 10.1126/SCIIMMUNOL.AAI7793,.

38. Lledó-Delgado, A., Preston-Hurlburt, P., Currie, S., Clark, P., Linsley, P.S., Long, S.A., Liu, C., Koroleva, G., Martins, A.J., Tsang, J.S., et al. (2024). Teplizumab induces persistent changes in the antigen-specific repertoire in individuals at risk for type 1 diabetes. Journal of Clinical Investigation 134. 10.1172/JCI177492,.

39. Baas, M., Besançon, A., Goncalves, T., Valette, F., Yagita, H., Sawitzki, B., Volk, H.D., Waeckel-Enée, E., Rocha, B., Chatenoud, L., et al. (2016). TGFβ-dependent expression of PD-1 and PD-L1 controls CD8+ T cell anergy in transplant tolerance. Elife 5. 10.7554/ELIFE.08133,.

40. You, S. (2015). Differential sensitivity of regulatory and effector T cells to cell death: A prerequisite for transplant tolerance. Front. Immunol. 6. 10.3389/FIMMU.2015.00242,.

41. Chatenoud, L., Thervet, E., Primo, J., and Bach, J.F. (1994). Anti-CD3 antibody induces long-term remission of overt autoimmunity in nonobese diabetic mice. Proc. Natl. Acad. Sci. U. S. A. 91, 123–127. 10.1073/PNAS.91.1.123,.

42. Kohm, A.P., Williams, J.S., Bickford, A.L., McMahon, J.S., Chatenoud, L., Bach, J.-F., Bluestone, J.A., and Miller, S.D. (2005). Treatment with Nonmitogenic Anti-CD3 Monoclonal Antibody Induces CD4+ T Cell Unresponsiveness and Functional Reversal of Established Experimental Autoimmune Encephalomyelitis. The Journal of Immunology 174, 4525–4534. 10.4049/JIMMUNOL.174.8.4525,.

43. Ota, T., Goto, R., Harada, T., Forgioni, A., Kanazawa, R., Ganchiku, Y., Kawamura, N., Watanabe, M., Fukai, M., Shimamura, T., et al. (2024). TCF1highPD-1+Ly108+CD8+ T Cells Are Associated with Graft Preservation in Sensitized Mice Treated with Non-Fc Receptor-Binding CD3 Antibodies. Immunohorizons 8, 295–306. 10.4049/IMMUNOHORIZONS.2300117.

44. Gavin, M.A., Clarke, S.R., Negrou, E., Gallegos, A., and Rudensky, A. (2002). Homeostasis and anergy of CD4+CD25+ suppressor T cells in vivo. Nat. Immunol. 3, 33–41. 10.1038/NI743,.

45. Levine, A.G., Arvey, A., Jin, W., and Rudensky, A.Y. (2014). Continuous requirement for the TCR in regulatory T cell function. Nat. Immunol. 15, 1070– 1078. 10.1038/NI.3004,.

46. Jing, Y., Kong, Y., Allard, D., Liu, B., Kolawole, E., Sprouse, M., Evavold, B., Bettini, M., and Bettini, M. (2023). Increased TCR signaling in regulatory T cells is disengaged from TCR affinity. bioRxiv. 10.1101/2023.01.17.523999.

47. Zhang, R., Huynh, A., Whitcher, G., Chang, J.H., Maltzman, J.S., and Turka, L.A. (2013). An obligate cell-intrinsic function for CD28 in Tregs. Journal of Clinical Investigation 123, 580–593. 10.1172/JCI65013,.

48. DuPage, M., Chopra, G., Quiros, J., Rosenthal, W.L., Morar, M.M., Holohan, D., Zhang, R., Turka, L., Marson, A., and Bluestone, J.A. (2015). The chromatin-modifying enzyme Ezh2 is critical for the maintenance of regulatory T cell identity after activation. Immunity 42, 227–238. 10.1016/j.immuni.2015.01.007.

49. Philip, L.P.B., Schiffer-Mannioui, C., Le Clerre, D., Chion-Sotinel, I., Derniame, S., Potrel, P., Bas, C., Lemaire, L., Galetto, R., Lebuhotel, C., et al. (2015). Multiplex genome-edited T-cell manufacturing platform for “off-the-shelf” adoptive T-cell immunotherapies. Cancer Res. 75, 3853–3864. 10.1158/0008-5472.CAN-14-3321,.

50. Wang, X., Wu, X., Tan, B., Zhu, L., Zhang, Y., Lin, L., Xiao, Y., Sun, A., Wan, X., Liu, S., et al. (2024). Allogeneic CD19-targeted CAR-T therapy in patients with severe myositis and systemic sclerosis. Cell 187, 4890–4904.e9. 10.1016/j.cell.2024.06.027.

51. Chiesa, R., Georgiadis, C., Syed, F., Zhan, H., Etuk, A., Gkazi, S.A., Preece, R., Ottaviano, G., Braybrook, T., Chu, J., et al. (2023). Base-Edited CAR7 T Cells for Relapsed T-Cell Acute Lymphoblastic Leukemia. New England Journal of Medicine 389, 899–910. 10.1056/NEJMOA2300709,.

52. Oldenhove, G., Bouladoux, N., Wohlfert, E.A., Hall, J.A., Chou, D., Dos santos, L., O’Brien, S., Blank, R., Lamb, E., Natarajan, S., et al. (2009). Decrease of Foxp3+ Treg Cell Number and Acquisition of Effector Cell Phenotype during Lethal Infection. Immunity 31, 772–786. 10.1016/j.immuni.2009.10.001.

53. Zhou, X., Bailey-Bucktrout, S.L., Jeker, L.T., Penaranda, C., Martínez-Llordella, M., Ashby, M., Nakayama, M., Rosenthal, W., and Bluestone, J.A. (2009). Instability of the transcription factor Foxp3 leads to the generation of pathogenic memory T cells in vivo. Nat. Immunol. 10, 1000–1007. 10.1038/NI.1774,.

54. Robert, J., Feuillolay, M., de Temple-Llavero, M., Akossi, R.F., Mhanna, V., Cheraï, M., Fourcade, G., Charlotte, F., Tchitchek, N., Mi, T., et al. (2025). Expression of an interleukin-2 partial agonist enhances regulatory T cell persistence and efficacy in mouse autoimmune models. Nature Communications 16. 10.1038/S41467-025-60082-Z,.

55. Qin, S., Cobbold, S.P., Pope, H., Elliott, J., Kioussis, D., Davies, J., and Waldmann, H. (1993). “Infectious” transplantation tolerance. Science 259, 974–977. 10.1126/SCIENCE.8094901.

56. Wardell, C.M., Fung, V.C.W., Chen, E., Haque, M., Tan, D.F.H., Leca, M., Gillies, J., Spanier, J.A., Mojibian, M., Fife, B.T., et al. (2025). CAR Treg cells mediate linked suppression and infectious tolerance in islet transplantation in mice. Sci. Transl. Med. 17. 10.1126/SCITRANSLMED.ADP6519.

